# Dysregulation of Hepatitis B Virus Nucleocapsid Assembly with RNA-directed Small Ligands

**DOI:** 10.1101/2021.08.10.455820

**Authors:** Nikesh Patel, Fardokht Abulwerdi, Farzad Fatehi, Iain Manfield, Stuart Le Grice, John S. Schneekloth, Reidun Twarock, Peter G. Stockley

## Abstract

RNA sequences/motifs dispersed across the pre-genomic copy of the Hepatitis B Virus genome regulate formation of nucleocapsids *in vitro* in an epsilon/polymerase independent fashion. These multiple RNA Packaging Signals (PSs) form stem-loops presenting in each loop a core protein recognition motif, -RGAG-. Small, drug-like molecules binding these motifs were identified by screening an immobilized library with a fluorescently-labelled RNA oligonucleotide encompassing the most conserved of these sites. This identified 66 “hits”, with affinities ranging from low nanomolar to high micromolar in SPR assays. High affinity ligand binding is dependent on the presence of the -RGAG-motif, which also appears to be the common element in cross-binding to other PS sites. Some of these compounds are potent inhibitors of *in vitro* core protein assembly around the HBV pre-genome. Mathematical modelling confirms the potential of these novel anti-viral drug targets for disrupting replication of this major human pathogen. Preliminary structure-activity relationships of the highest affinity compound reveal critical functional groups for PS-binding. PS-regulated assembly is easily adapted to high-throughput screening allowing future development of pharmacologically active compounds.

## Introduction

Hepatitis B Virus (HBV) is estimated to have infected over 2 billion individuals and is the largest cause of liver cancer worldwide. Its heavy annual death toll and severe economic costs arise within the ∼260 million people who suffer from chronic infections^1^, for which there is currently no cure. These patients typically suffer cycles of asymptomatic liver inflammation, which over decades can lead to hepatocellular carcinoma and cirrhosis, and hence death. Despite the availability of an effective vaccine^2^, it is not universally deployed, allowing ∼1 million new infections annually, many of which are due to vertical transmission from mother to child, a route that frequently progresses to the chronic state. Currently available therapies for HBV are limited in scope. Interferon therapy is used as an adjuvant for the immune system but has had limited success in reducing HBV levels, and produces undesirable side effects^3,4^. Nucleot(s)ide analogues directed at the polymerase are more widely used, but require lifelong treatment, and can elicit resistance mutations.

HBV is a para-retrovirus, initially encapsidating a positive-sense, single-stranded (ss) ∼3500 nt RNA, termed the pre-genomic RNA (pgRNA), into a *T*=4 nucleocapsid (NC)^5^. This is widely believed to require co-assembly with a molecule of the virally-encoded polymerase which binds specifically to a site, the ε stem-loop on the pgRNA^6–9^. This enzyme reverse transcribes the pgRNA, whilst degrading it, and then copies the ssDNA strand formed, all within the confines of the NC^10–12^. These steps result in formation of the relaxed, circular dsDNA genome (rcDNA) found within mature HBV virions. Once within a host cell, rcDNA is repaired by host enzymes generating a covalently-closed circular DNA (cccDNA)^13,14^, Fig 1, from which the sub-viral mRNAs for expression of HBV proteins are transcribed^5,9^. Given the initial requirement to package a ssRNA genome, we explored previously whether this virus assembles its NC using the RNA Packaging Signal (PS)-mediated mechanism we have identified in the genomes of many spherical ssRNA viruses from other viral families^15–21^ These encompass multiple, dispersed sequences, often in the form of stem-loops, that act to regulate assembly of the infectious virion^15–17,19,20,22^. Within a viral gRNA PS sites share affinity for their cognate coat proteins (CPs) resulting in improved assembly efficiency and sequence-specificity, whilst ensuring genetic resilience in a group of pathogens that replicate via error-prone polymerases^23^. Recently, we showed that an analogous mechanism evolves spontaneously in an artificial self-assembling bacterial enzyme^18^, implying that its utility extends beyond virology.

**Figure 1:**
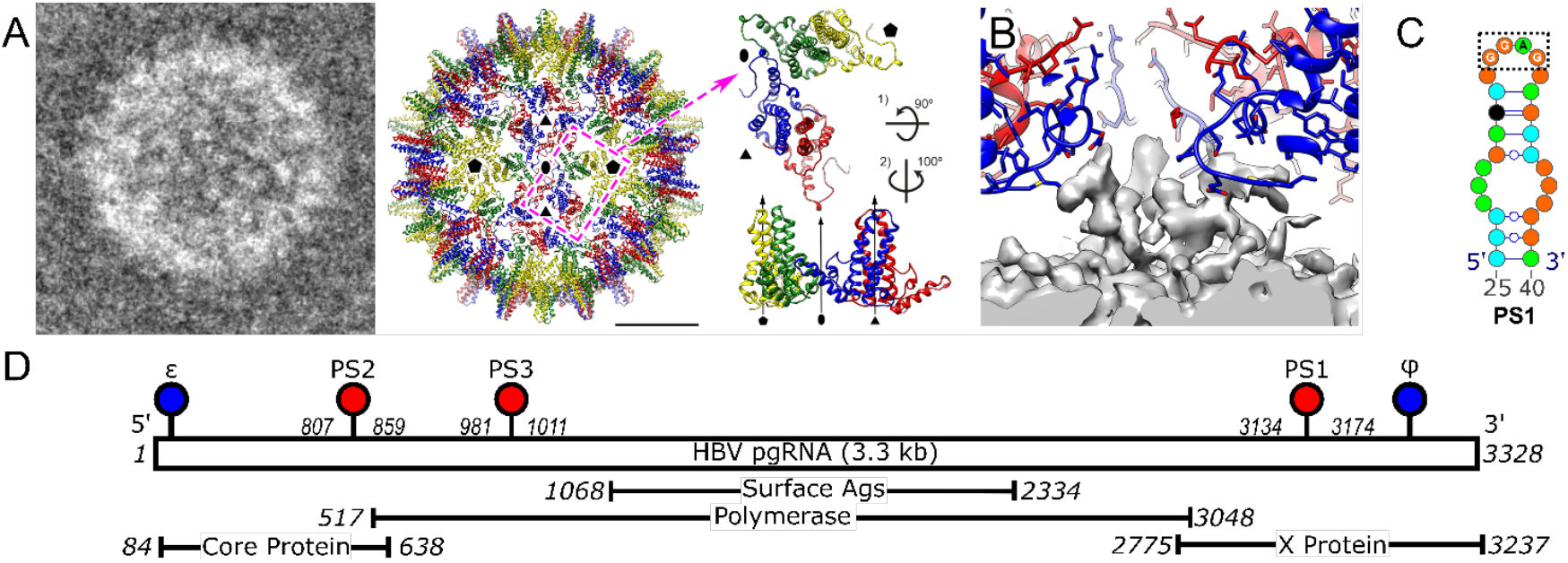
Packaging Signals trigger nucleocapsid reassembly. (A) *Left*, negatively stained micrograph of a reassembled *T*=4 NC. *Centre*, Atomic model of the front-half capsid of HBV *T*=4 NC shown as ribbon diagrams, coloured in yellow (A monomer), green (B monomer), blue (C monomer) and red (D monomer) for the quasi-equivalent monomers, and viewed along a two-fold axis (PDB ID: 7ABL,^25^). The dashed magenta rectangle indicates the asymmetric unit composed of 2 Cp dimers shown, *right*. The 3-fold axes between C & D monomers are the locations of the Cp-pgRNA contacts shown in panel B. (B) The atomic model of Cp dimers around the 3-fold axis is fitted into the non-filtered density map of pgRNA-filled NC (EMD-11702). Contacts located at three-fold vertices involve the E40-C48 sequence in Cp (including the E40-C43 alpha-helix and the S44-C48 loop) of three C monomers. The side chains of residues E40 and E46 directly point at the pgRNA shell (C) The PS1 (NC003977.1) stem-loop used as the target for the SMM screen. G:C clamps were used to force the presentation of the bulged structure in the sequence from the NC_003977.1 strain, presenting the -RGAG-motif on an apical loop (Nucleotides are coloured as follows: Light green=A, orange=G, black=C, cyan=U). Watson-Crick base pairs for RNA oligomers are indicated as lines, interrupted by circles for G-U pairs. See also Supplementary Information. (D) Genetic map of the pgRNA of HBV, strain JQ707375.1. The positions of its ORFs with their nucleotide numbering are indicated below. RNA stem loops ε, ψ (blue), PS1, PS2 and PS3 (red), are indicated as lollipops above.

RNA SELEX^24^ against full-length Cp from strain NC_003977.1, yields aptamers with sequence matches to the underlying gRNA allowing identification of up to 16 HBV PSs within pgRNA sequences of >300 strains. The three most evolutionarily-conserved PS sites in strain NC_003977.1 (PSs1-3) each trigger sequence-specific assembly of Cp dimers *in vitro* into mostly *T*=4 NC-like particles (NCPs)^19^. Cryo-EM of these NCPs suggests that the PSs are sequence-specific triggers of in vitro assembly. Consistent with this idea, homologues of these PSs in the pgRNA of a distinct HBV strain (JQ707375.1) regulate nuclease-resistant assembly of *T*=4 NCPs with the Cp from NC_003977.1 (Fig 1A, B). The homologues also form stem-loops, containing the expected -RGAG-motif (Fig. 1C). Each PS contributes individually and collectively to the efficiency of NCP formation *in vitro*^25^. Since the PS-mediated assembly mechanism appears to regulate NC formation, we used a small molecule microarray (SMM)^26–30^ to identify sequence-specific binding ligands to an oligonucleotide encompassing PS1. Thereby, we identified multiple ligands that as non-immobilised versions bind tightly to oligonucleotides encompassing PS1, and other PS sites. The highest affinity compounds are potent inhibitors of NCP formation in the presence of the JQ707375.1 pgRNA. Preliminary structure-activity analysis of the best inhibitors reveals the critical features of the inhibitory ligands, whilst high-throughput plate-based assays suggest that this system offers a novel route for identification of HBV inhibitors. Mathematical modelling confirms the therapeutic potential of such directly-acting anti-virals (DAA)^36^.

## Methods

### Cloning, expression and purification of HBV Cp dimer

HBV Cp was expressed from a pET28b plasmid in *E. coli* BL21(DE3) cells and purified as NCPs ^19,31^. NCPs were dissociated into Cp dimers using 1.5 M guanidinium chloride as previously described^19,31^. The HBV Cp dimer concentration was determined spectrophotometrically (ε_280_ of Cp dimer = 55,920 L mol^-1^ cm^-1^) in a Nanodrop™ One. Fractions with an A_260/280_ ratio of ∼0.65 or lower were used in reassembly assays. Absorbance values at 260 and 280 nm were corrected for light-scattering throughout, using the absorbance values at 310 and 340 nm, as previously described^32^.

### Preparation of pregenomic RNA

A wild-type pgRNA construct was assembled using a clone purchased from ATCC® (pAM6 39630™, strain acc. no JQ707375.1). The pgRNA precursor sequence was copied in a modular fashion using PCR and cloned into the correct order within a pACYC184 vector, between the *BspHI* and *HindIII* sites using a Gibson reassembly® Master Mix, according to the manufacturer’s protocol (New England Biolabs). Transcription of the RNA was carried out using a Highscribe T7 High-yield RNA synthesis kit (NEB), after linearisation of the DNA plasmid using *HindIII*. RNA was annealed by heating to 70 °C for 90 sec followed by cooling slowly to 4 °C in a buffer containing 50 mM NaCl, 10 mM HEPES and 1 mM DTT at pH 7. Products were assessed using a 1% (w/v) denaturing formaldehyde agarose gel. RNA concentration was estimated using A_260_ values (ε_260_ of pgRNA = 32, 249 mM^-1^ cm^-1^).

### Small molecule microarray screening

SMM screening was performed as previously described^26^. Briefly, Corning GAPS II slides were functionalized with an isocyanate group using an established procedure^33^. Slides were printed using an ArrayIt NanoPrint LM60 microarrayer, and compounds were printed as 10 mM stock solutions in DMSO with two replicate spots. After printing, slides were incubated with fluorescently-labelled PS1 RNA purchased from Dharmacon. After washing, slides were imaged with an Innopsys 1100 scanner. Composite Z scores were generated for each compound. Hits were characterized as (1) having a Z score >3, having an increase of Z-score of 3 relative to a buffer-incubated control, and (3) having a coefficient of variance (CV) of <120% for replicate spots. Finally, hits were filtered for selectivity against all other RNAs screened previously in the SMM format in the Schneekloth lab.

### Assaying ligand affinities for PS RNAs

#### RNA preparation

5’-amino-labelled RNA oligonucleotides (Table 1) were purchased from IDT and biotinylated covalently by rolling at room temperature for 4 h in 100 mM sodium borate buffer (pH 8.0), with a 100-fold molar excess of EZ-Link Sulfo-NHS-LC-LC-Biotin (Thermo Fisher). Biotinylated oligonucleotides were extracted from a 10% (v/v) denaturing acrylamide gel via a ‘crush and soak’ method. Gel slices corresponding to the desired oligonucleotides were excised and eluted three times into separate aliquots (100 μL) of 10 mM TE buffer over 3 h. Aliquots were combined and precipitated overnight using 50%(v/v) isopropanol, 0.3 M NaOAc and 1/100 volume RNA grade glycogen at -20°C. Biotinylated, precipitated RNA was eluted into 100 μL ddH_2_O.

**Table 1.**
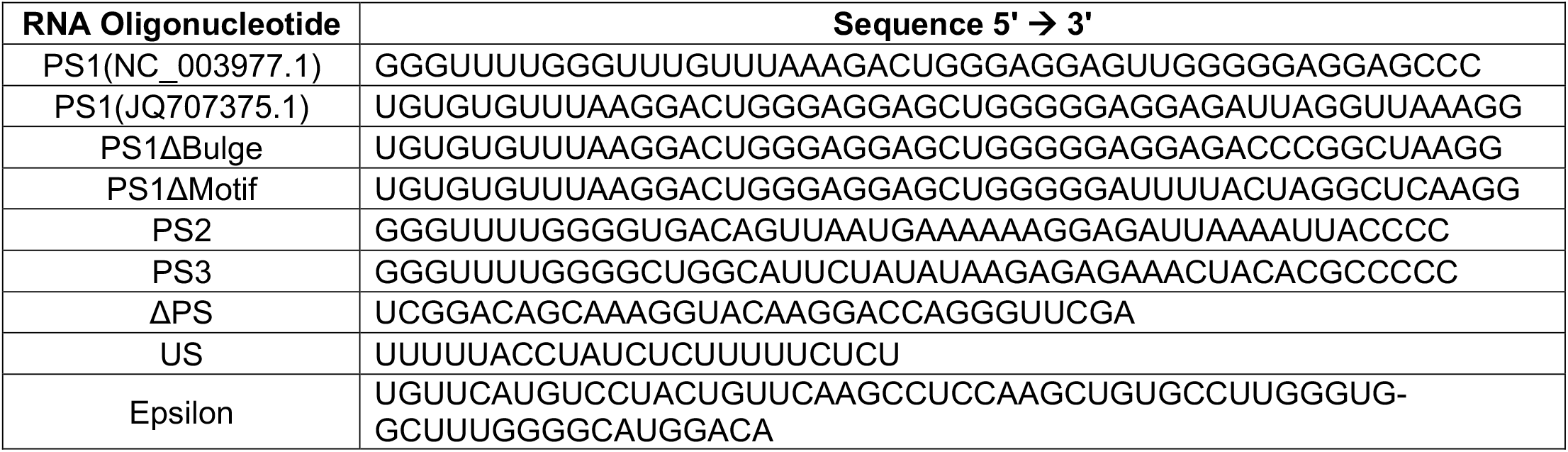
RNA Oligonucleotides used in SPR binding assays.

#### Determining ligand affinities using surface plasmon resonance (SPR)

Biotinylated RNA oligonucleotides were immobilised onto the surface of streptavidin-coated chips (GE Healthcare) in a BIAcore T200 at a concentration of 100 nM. Buffer flow was set at 5 µL/min over 15 min contact time, and the surface of the chip saturated with the RNA oligonucleotides, as determined by a plateau in the resulting refractive index change. Small molecular weight compounds stored in DMSO at a concentration of 10 mM, were used at concentrations of 75, 100, 150, 200 and 300 µM. These were stored in a chilled compartment at 9°C within a skirted 384 well plate (GE Healthcare) before being washed over the chip surface in a buffer containing 20 mM HEPES (pH 7.5), 250 mM NaCl, 2%(v/v) DMSO and 0.1%(v/v) Tween20, at a rate of 10 µL/min for 6 min, allowing association of 1 min followed by dissociation of 5 min. Solvent corrections to account for variable concentrations of DMSO were performed according to manufacturer’s instructions. Data were plotted with the best-fitting model as determined by the chi^2^ values. All fits used the two-state model available in the Biacore T200 evaluation software, detailed below (GE Healthcare).

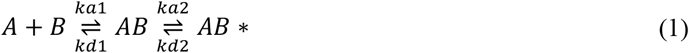

The two-state model (1) used to fit the SPR curves. The association (*k*_*a1*_, *k*_*a2*_) and dissociation (*k*_*d1*_, *k*_*d2*_) rate constants of complexes AB and AB*, where A is the analyte (compound) and B is the immobilised RNA, were determined.

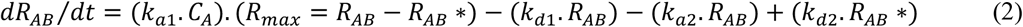

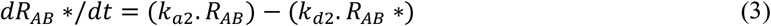

Equation (3) defines the rate of formation of each complex, AB (2) and AB* (3), directly from the SPR response curves obtained. R is the concentration of the complex (subscript) and C_A_ is the concentration of the analyte.

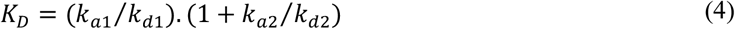

Dissociation and association constants from (2) and (3) were used to calculate an affinity constant (4)^34,35^.

#### HBV NCP Reassembly

178 μL of 1.1 nM heat-annealed pgRNA in a buffer containing 20 mM HEPES (pH 7.5), 250 mM NaCl and 5 mM DTT, was incubated in each well of a 96 well plate at room temperature for 30 min. 2 μL of DMSO ± compound was added, and a further equilibration was performed for 30 min. Cp dimer in dissociation buffer (as above) was then titrated into the RNA using a Biomek 4000 liquid-handling robot (Beckmann Coulter), step-wise up to a ratio of 1200:1 (10 discrete 2 μL solution aliquots, using the following Cp dimer concentrations (in brackets), were then added: 1 (100 nM), 10 (1 μM), 25 (2.5 μM), 75 (7.5 μM), 120 (12 μM), 240 (12 μM), 480 (24 μM), 720 (24 μM), 960 (24 μM) and 1200 nM (24 μM) Cp dimer). The final volume in each well was 200 μL, i.e. a cumulative volume of 19.2 mL/plate. The final concentrations of RNA and Cp were 1 nM and 1.2 μM, respectively. Cp aliquots were calculated such that they reached a maximum of 10% of the final volume, maintaining the GuHCl concentration <0.15 M. Following incubation at room temperature for 1 h, the samples were pooled and concentrated to a final volume of 2 mL. The A_260/280_ ratio was measured using a Nanodrop™ One (Thermofisher Scientific), and the RNA concentrations calculated using the corrected absorbance value at 260 nm. Each sample was split into 2, with one half treated with 1 µM RNase A, and incubated overnight at 4 °C. After incubation, 5 µL of each sample was visualised by negative stain electron microscopy (nsEM) to assess particle yield, shape and “completeness”. The remaining samples were analysed by application to a TSK G6000 PWXL column (Tosoh), in a buffer containing 20 mM HEPES (pH 7.5), 250 mM NaCl and 5 mM DTT, attached to a SEC-MALLS system (ÄKTA Pure (GE Heathcare) connected to an Optilab T-REX refractometer and miniDAWN^®^ multiple angle laser light-scatterer fitted with a Wyatt QELS DLS module (Wyatt Technology)). Light-scattering peaks were collected and concentrated to ∼1 ml, where their RNA content, was estimated using their A_260_ relative to the starting material.

#### Mathematical modelling of the effects of ligands on NC assembly

Assembly kinetics were modelled via a set of reactions adapted from our recent intracellular model for HBV infection^36^, that includes the roles of the three evolutionarily conserved packaging signals (PS1-PS3) in the pgRNA in virion assembly^19^. We assume that all three PSs interact with Cp dimers and PS-targeting compounds according to the following reactions:

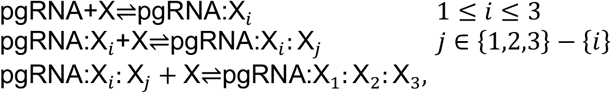

where X denotes either a Cp dimer (C) or a compound (d), and an index refers to the PS site where binding occurs. For example, pgRNA:C_1_: C_2_: d_3_indicates C bound to PSs 1 and 2, and a drug molecule to PS3. The pgRNA:C_1_: C_2_: C_3_complex will then recruit 117 Cp dimers at a rate κ to build an intact particle via recruitment of further CP dimers:

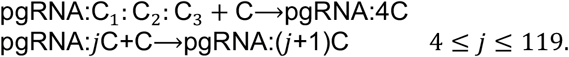

In order to model malformations caused by drug molecules, we introduce reactions in which the number of Cp dimers to be recruited is limited to *n*_*c*_. The latter is estimated based on the experimentally determined hydrodynamic radii (*R*_*h*_) of the malformed particles (Tables 2 & 3). In particular, in the compound-free case we have *R*_*h*_ = 18.2 nm and each nucleocapsid contains 120 Cp dimers. As the surface of a sphere is equal to 4π*r*^2^, we assume that the number of Cp dimers is in a good approximation proportional to 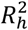. Therefore, we use 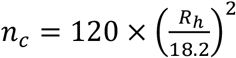 for the average number of Cp dimers in a malformed particle. For example, *n*_c_ ≈ 114 for *R*_*h*_ = 17.7 nm.

**Table 2:** Binding affinities (nM) and associated standard error of the mean (errors <1 are omitted) of PS binding compounds characterised in this study, for RNA oligonucleotides encompassing PS1 from different strains of HBV (Left to right, NC_003977.1, JQ707375.1) and PS3. Full table available in Supplementary Information (Sup Table 1)

| Compound | $K_D$ / nM | | |
| --- | --- | --- | --- |
|  | PS1(NC_003977.1) | PS1 (JQ707375.1) | PS3 |
| 3 | 211982 ± 1179 | 224056 ± 455 | 99061 ± 591 |
| 10 | 11774 ± 284 | 11374 ± 170 | 8539 ± 797 |
| 46 | 10115 ± 119 | 10196 ± 225 | 32658 ± 227 |
| 63 | 12 | 19 | 15 |

#### Parameter value estimation

178 μl of 1.1 nM pgRNA in the drug-free scenario corresponds to 1.179 × 10^11^ copies of pgRNA. The simulation mimics the experimental protocol for Cp dimer titration with step-wise addition in 10 discrete steps with 10 minutes intervals up to a ratio of 1200:1. The numbers of Cp dimers added at consecutive steps were; 1.2 × 10^11^, 9 × 1.2 × 10^11^, 15 × 1.2 × 10^11^, 50 × 1.2 × 10^11^, 45 × 1.2 × 10^11^, 120 × 1.2 × 10^11^, 240 × 1.2 × 10^11^, 240 × 1.2 × 10^11^, 240 × 1.2 × 10^11^, and 240 × 1.2 × 10^11^. After the equivalent of one hour incubation time, assembly reactions are started with recruitment rate κ equal to 10^6^ M^-1^S^-1 37^ and PS binding affinity for Cp of 4 nM ^36^.

As 1 OD_260_ Unit = 40 μg/ml and assuming a volume of 100 *μ*l and given the average weight of a nucleocapsid of 4 MDa (6.6422 × 10^−18^g) ^38^, we obtain 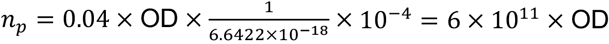 for conversion of OD values to the number of assembled particles (*n*_5_). Since after nuclease treatment (160 minutes post experiment start) A260 is 0.12 (Table 3), the number of fully formed nucleocapsids are equal to 7.2 × 10^19^. We fitted the forward rate of Cp dimers for binding to PSs (*f*_*n*uc_) to match this value, giving us 336 M^−1^S^−1^.

**Table 3:** Effect of Compounds on pgRNA mediated NCP reassembly in vitro. Left to Right: A_260_ (before and after nuclease treatment), A_260/280_ ratios, packaged RNA yield, and R_*h*_ values for reassembled materials resulting from the interaction between Cp and pgRNA, in the absence and presence of #3, 10, 46 and 63. Values are shown after treatment with RNase A.

| Compound | NCP Peak |  |  |  |  |
| --- | --- | --- | --- | --- | --- |
|  | A<sub>260</sub> (Before- → After Nuclease treatment) | A<sub>260/280</sub> | RNA yield | R<sub>h</sub> / nm | NCP appearance |
| N/A | 0.13 → 0.12 | 1.43 | 92% | 18.2 | Separate, stain excluding NCPs |
| 3 | 0.021 → 0.014 | 1.42 | 67% | 18.0 | Stain Excluding NCPs with single “bleb” |
| 10 | 0.027 → 0.017 | 1.23 | 63% | 15.2 | Mostly large “flattened” NCPs |
| 46 | 0.009 → 0.003 | 0.76 | 33% | 14.7 | Mostly large “flattened” NCPs |
| 63 | 0.025 → 0.004 | 0.98 | 16% | 17.2 | Aggregated partially formed particles |

In the presence of 1 nM of compound 63, A_260_ is 0.028 after nuclease treatment. Using the formula above, *n*_*p*_ ≈ 1.7 × 10^10^. We explored different forward rates for the binding of compound 63 to PSs (*k*_*f*_). For *k*_*f*_ < 5 *μ*M^−1^S^−1^, the number of fully formed particles is > 1.7 × 10^10^. Increasing *k*_*f*_ from 5 to 10 *μ*M^−1^S^−1^ reduces the number of nucleocapsids to 1.4 × 10^10^. However, further increasing of *k*_*f*_ does not reduce the number of nucleocapsids and it stays around 1.4 × 10^10^ particles. We therefore show results for *k*_*f*_ = 10 *μ*M^−1^S^−1^, and also use this value for other compounds, which is consistent with an estimate of 1 to 10 *μ*M^−1^S^−1^ for other molecules^39^.

#### Fluorescence anisotropy NCP assembly assay

178 μL of 1.1 nM heat-annealed pgRNA or 16.5 nM heat-anneal PS1(NC_003977.1) in a buffer of 20 mM HEPES (pH 7.5), 250 mM NaCl and 5 mM DTT, was incubated in wells of a 96 well plate at room temperature for 30 min. Each nucleic acid was fluorescently labelled at the 5′ end using an Alexa Fluor SDP ester as previously described^19^. 2 μL of DMSO ± compound was added, and a further equilibration was performed for 30 min. Cp dimer in dissociation buffer (as above) was then titrated into the RNA using a Biomek 4000 liquid-handling robot (Beckmann Coulter), step-wise up to a ratio of 1200:1 (10 discrete 2 μL solution aliquots, using the following Cp dimer concentrations (in brackets), were then added: 1 (100 nM), 10 (1 μM), 25 (2.5 μM), 75 (7.5 μM), 120 (12 μM), 240 (12 μM), 480 (24 μM), 720 (24 μM), 960 (24 μM) and 1200 nM (24 μM) Cp dimer). RNA only controls were performed, with dissociation buffer added without Cp dimer. The final volume in each well was 200 μL. Fluorescence anisotropy was measured after each addition using a POLARstar OMEGA (BMG Labtech, Aylesbury). The final concentrations of RNA and Cp were 1 nM and 1.2 μM, respectively. Cp aliquots were calculated such that they reached a maximum of 10% of the final volume, which keeps the GuHCl concentration <0.15 M. Each well was then treated with 1 µM RNase A, and incubated at room temperature for 1 hour, after which a final fluorescence anisotropy reading was taken. These values were normalised with respect to the initial reading, and the anisotropy change calculated as the final RNase treated sample – the starting fluorescence anisotropy of the RNA.

## Results

### Identification & Characterisation of PS1 Binding Compounds

The 47-nt HBV packaging motif labeled with the red fluorescent dye (AlexaFluor 647) was used for small molecule microarray screening. A library of 20,000 drug-like molecules was covalently linked onto an isocyanate-functionalized glass surface. Slides were incubated with 70µL of packaging RNA (5µM) while a second copy of the same slide was incubated with the buffer used to anneal the RNA. A control buffer-incubated slide eliminated selection of any auto-fluorescent compounds. After two washes, slides were scanned at 635nm and statistical analysis was performed. 66 ligands that bound the novel HBV packaging signal were thus identified.

### PS-Ligand Affinities

Commercially-available PS1-binding compounds, identified from the SMM screening and numbered by their order in that array, were purchased and resuspended in DMSO (10 mM) and stored frozen until diluted for binding assays. SPR was used to screen each ligand for binding affinity for a series of RNA oligonucleotides. These included PS1 (from both NC_003977.1 the initial aptamer and drug selection target; & JQ707375.1, Fig 1C); and sequence/structure variants, including the NC_003977.1 PS2 & PS3, bulge & loop variants of the NC_003977.1 PS1 (PSΔBulge and PSΔMotif, respectively), and an unrelated stem-loop (ΔPS) lacking both the motif and the single-stranded bulge (Fig 2, Sup Fig 2). RNAs were biotinylated at their 5′ ends and immobilised onto sensorchips coated with streptavidin. The affinities and stoichiometries of the test compounds for these sites were then determined from the apparent association (*k*_*a*_) and dissociation (*k*_*d*_) rate constants of the complexes formed (Methods). The best fits to the data used the two-state binding model present within the T200 Biacore Evaluation Software (Table 2, Fig 2,Sup Table 1). Most compounds possessed higher affinities for the PS1 oligonucleotide they were initially elected against than the other RNAs tested. Roughly a third have significantly better binding to HBV PSs other than PS1. The latter result is consistent with the functional conservation of the PSs despite sequence/structural changes. The worst binders were compounds 27, 54 and 55 which have affinities in the millimolar range. All compounds bound preferentially to PSs 2 and 3 and negative controls, implying recognition of a common feature. Only 5 of the 66 compounds tested show any binding to the controls with affinities in the high millimolar range (compounds 1, 4, 25, 49 and 50, data not shown). Compounds affinities (*K*_*D*_) range from nanomolar to millimolar. The RU changes on binding suggest a stoichiometry of 2:1 ligand: PS RNA for the highest affinity binders, consistent with the model applied to analyze the data. Lower affinity binders appear to bind less specifically with an apparent saturations ∼5:1 ligand: PS RNA.

**Figure 2:**
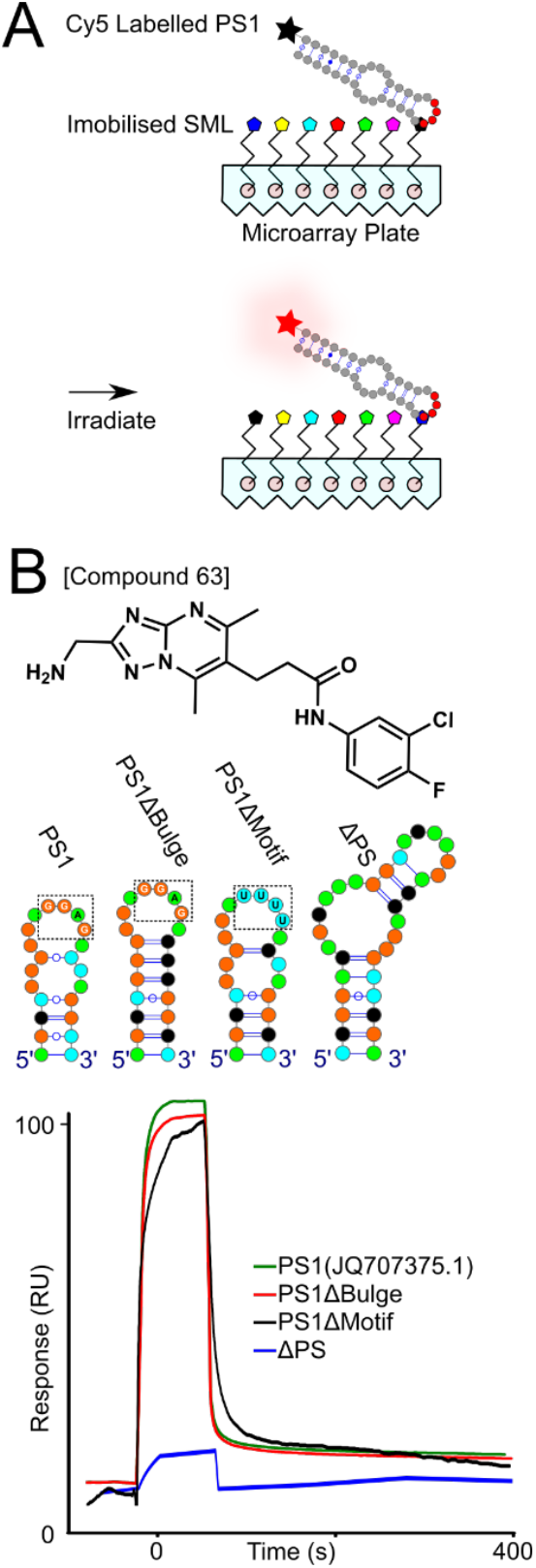
PS RNA binding compounds are identified using a SMM and their affinities measured using SPR. (A) SMM assay identifying PS1 RNA binding compounds. Cy5 (black star) labelled PS1 was washed over immobilised small molecular weight ligands (pentagons), with binders identified by the presence of fluorescence (red star) after irradiation. Full SMM screen available in Sup Fig 1. (B) *Top*, Structure of #63 tested in SPR traces shown below. *Middle*, PS1, PS1ΔBulge, PS1ΔMotif and ΔPS stem loops. (Light green=A, orange=G, black=C, cyan=U). Free folding energy and full structures for RNA oligomers are shown in Sup Fig 2. Watson-Crick base pairs for RNA oligomers are indicated as lines, which are interrupted by circles for G-U pairs. *Bottom*, SPR traces for interactions between 75 µM compound 63 and PS1 (green), PS1ΔBulge (red), PS1ΔMotif (black) and ΔPS (blue).

### Inhibition of NCP Assembly

In order to test the whether the PS-binding ligands identified would inhibit assembly of the HBV NCP *in vitro*, we established assays at low (nanomolar) gRNA concentrations. We have shown previously in single molecule fluorescence assays that assembly under these conditions is dependent on sequence-specific interactions between Cp and gRNA^19,25^, i.e. it at least mimics *in vivo* assembly outcomes^40^. The titration of Cp into pgRNA was automated on a liquid-handling robot (Methods). This allows preparation of reassembled material in 96-well plates at low concentrations, all wells subsequently being pooled and concentrated. Here we used a ∼3200 nt pgRNA (strain JQ707375.1), originally isolated from a patient in a lamivudine resistance mutation study^41^. This construct possesses the ε stem loop and structure/sequence homologues to the previous PSs^25^. It lacks the additional ∼300 nts in the pgRNA that would be present within a *bona fide* infection. Also absent is the 5′ cap and poly-A tail. As the viral polymerase is not easy to prepare in a functional form for *in vitro* study, this construct monitors NCP formation regulated only by the PSs^25^. It would therefore be sensitive to disruption of PS-Cp interactions.

In order to assess the effects of PS-binding ligands on assembly, small-scale reactions (90 μL) at nanomolar concentrations were performed, titrating 1.2 µM Cp in 10 steps over a 2h period into a solution of pgRNA (1.1 nM, final concentration 1 nM). Ligands were added (final concentration 10 μM) to the pgRNA solution prior to Cp titration, allowing the full range of ligand affinities to be tested. Pooled samples from 96-wells were then split into two aliquots, one of which was treated with RNase A to degrade any unencapsidated pgRNA. A sample of each of these was then visualised by nsEM (Fig 3). The remaining sample was fractionated by size-exclusion chromatography, monitoring the outflow via a multiple-angle, laser light-scattering detector (SEC-MALLS). These assays allow the integrity, shape, and hydrodynamic radius (R_*h*_) of assembled particles to be assessed whilst providing an estimate of assembly efficiency (Sup. Fig. 4). Peaks eluting from the gel filtration column were re-concentrated to determine their RNA and Cp content via their A_260/280_ ratios (Table 3). A typical uninhibited reaction in the presence of DMSO is shown in Sup. Fig. 4. There is a dominant, single symmetrical light-scattering peak (red) eluting from the column at ∼ 8-8.5 mLs post-injection. The peak fraction has an R_*h*_ of 18.2 nm, slightly smaller than that of NCPs produced by Cp expression in *E*.*coli* in the absence of pgRNA (19.4 nm, ^25^). Nuclease-treatment produces very little change in this outcome (green trace). EMs reveal discrete, stain-excluding, single particles of roughly the expected diameter. The fact that the particles exclude stain is consistent with their pgRNA content. Their A_260/280_ ratios are largely unaffected by nuclease treatment (Table 3) implying that the assembled Cp shells are complete. Note, the filter used to concentrate samples removes unassembled Cp from the traces.

**Figure 3:**
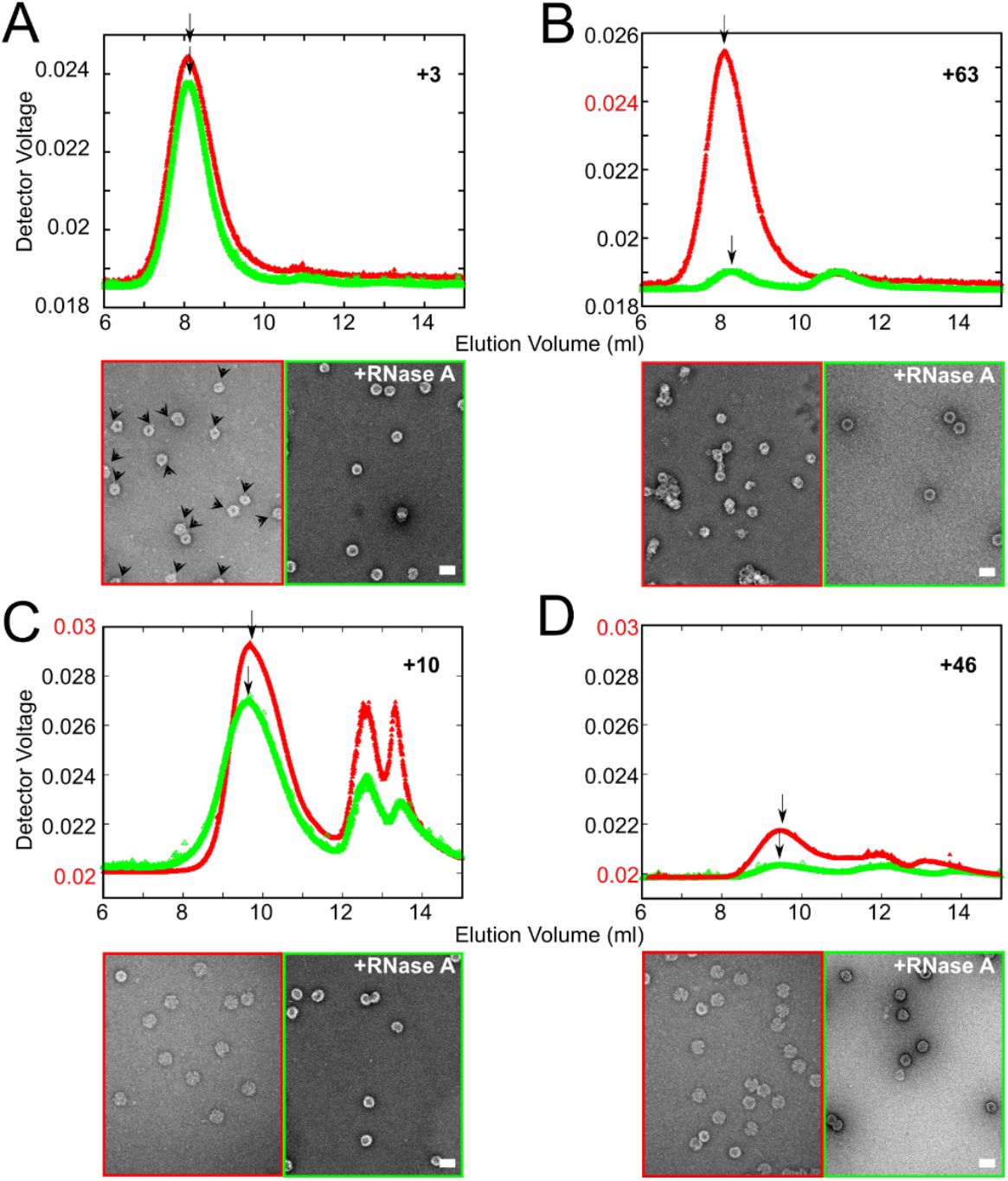
Reassembly assays to determine the effect of PS binding compounds on NCP reassembly. µM Cp dimer was titrated in gradually increasing increments into 1 nM heat annealed (Methods) pgRNA. These samples were characterized as previously described^25^, using a SEC-MALLs system and nsEM. (A) 1 nM pgRNA and Cp dimer in the presence of #3, (B) 1 nM pgRNA and Cp dimer in the presence of #63, (C) 1 nM pgRNA and Cp dimer in the presence of #10 and (D) 1 nM pgRNA and Cp dimer in the presence of #46 -(red)/+(green) nuclease treatment. R_*h*_ values were taken at the arrows above each peak. *Above*, light scattering (LS) traces of the reassembly mixes, coloured-coded as described above. *Lower*, nsEMs of reassembly mixes colour-coded as in LS traces. Scale bars throughout = 50 nm.

Compound 3, which has one of the weakest affinities for PS1 (*K*_*D*_ ∼ 210 µM), but has a ∼700-fold better binding to PS2 (*K*_*D*_ ∼ 300 nM), would not be expected to saturate PS3 and arguably the most important PS, PS1, at a concentration of 10 µM. Consistent with this, in the assembly assay it forms largely nuclease-resistant NCPs around pgRNA (Fig 3A) that elute in the same position as those lacking ligand. The A_260/280_ ratios of these peaks are also similar to the control, implying that Compound #3 has very little effect on the NCPs that do form. However, the NCP yield is reduced dramatically (>5-fold) compared to the control. Although batches of Cp can be variable depending on the amount of guanidinium ions present, this level of variation is striking (Tables 2 & 3). EM images offer some insights into this outcome. Prior to nuclease treatment, most reassembled particles lack the smooth circular appearance of the uninhibited control, and stain penetration is variable. In addition, many particles appear to have a unique structural feature on their peripheries that appears bright compared to the rest of the particle (see arrowheads on the micrograph, Fig 3A). These features seem mostly to disappear with nuclease treatment implying that they are due to pgRNA. Although the impact of Compound #3 appears limited, this hints that it is partially inhibiting correct encapsidation, consistent with the drop in NCP yield, which is potentially due to differential recovery from the concentration step.

Compound #63 has the highest affinity for PS1 (*K*_*D*_ ∼ 19 nM) and shows the strongest inhibitory effects in NCP assembly assays. The sample prior to nuclease treatment possesses the same apparent *R*_*h*_ as the control, with a yield similar to that obtained with Compound #3. However nuclease treatment destroys these particles yielding only minor background peaks. Strikingly, in the presence of Compound 63 particles viewed by nsEM are very different than the control. Post RNase treatment, most particles are heterogeneous, forming large clumps with relatively few remaining as separate shells of stain-penetrated particles of the correct size (Fig 3B). The absorption data confirm these interpretations (Table 3). Similar assembly inhibitory effects were observed with Compounds 10 and 46, both of which have intermediate affinities for PS sites (Table 2, Fig 3C & D). Both these compounds result in low yield, nuclease-sensitive particulate material that elutes later than the uninhibited control peak. The results in each case show multiple additional peaks, implying that the assembly process is dysregulated by treatment with PS-binding compounds.

This effect is dependent on the affinity of the ligand for RNA, e.g. Compound #63 (Fig 4; Table 4). Post-RNase treated material shows a dramatic decrease in the yield of fully reassembled NCP, even at 1 nM ligand concentration, i.e. well below the apparent affinity for PS1. It appears that at stoichiometric concentrations with gRNA but at very low levels of PS saturation, assuming that the affinity with an oligonucleotide is similar to that of the longer RNA, potent dysregulation occurs. These effects become more pronounced as ligand concentration is titrated in 10-fold increasing steps up to 10 μM. Reduced NCP yield is accompanied by significant changes in the appearance of the samples in nsEM. At the lowest ligand concentration (1 nM) particles are largely separate and circular, although mostly stain-penetrated with some small variations in apparent diameter. This regularity is lost with ligand titration, reflected by incomplete and malformed shells and increased clumping. The latter behavior is consistent with aggregates formed by multiple protein complexes binding across several pgRNAs.

**Figure 4:**
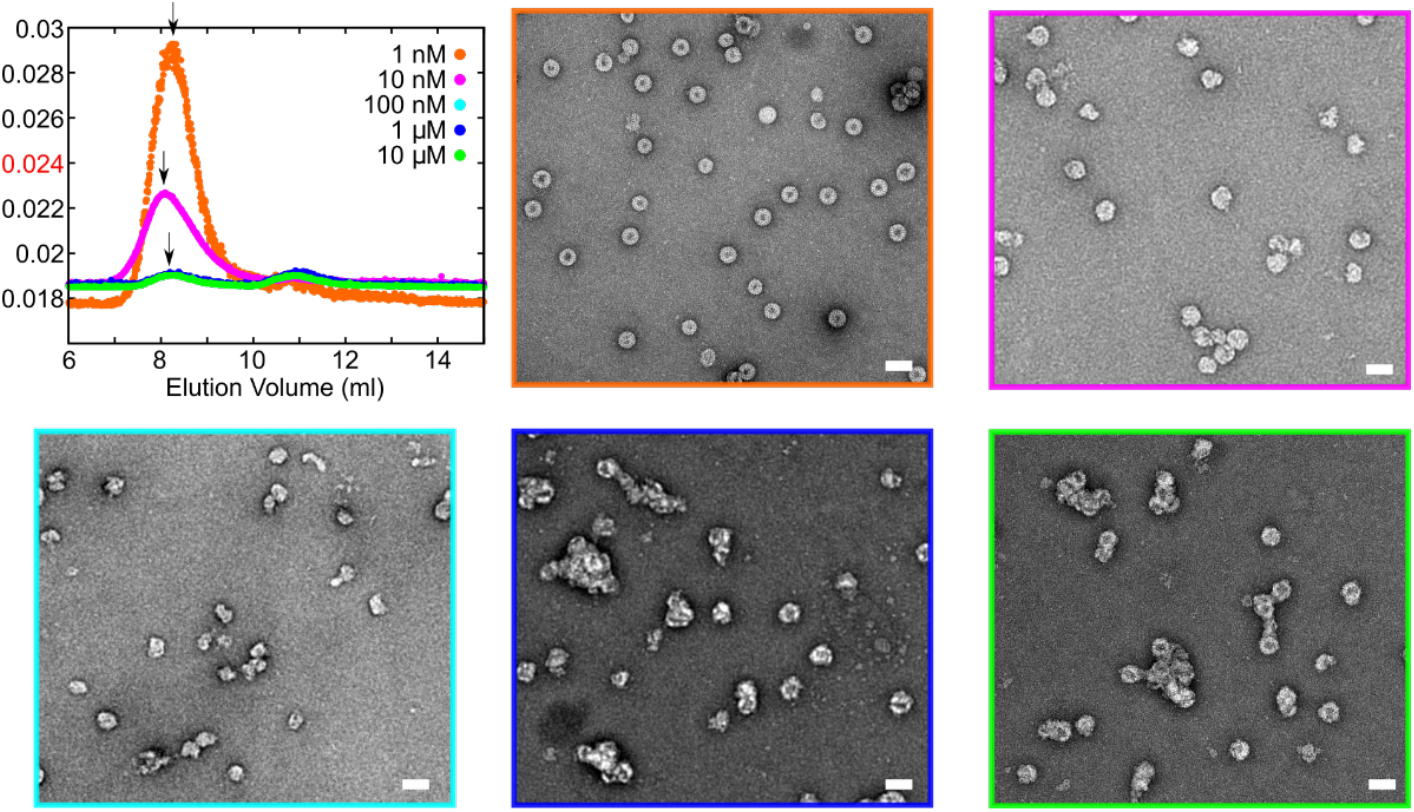
Investigating the limits of inhibition by compound 63. *Top left*, LS traces of the NCP reassemblies (performed as described in Methods and previously^25^) between 1nM pgRNA and Cp dimer in the presence of: 1 nM (orange), 10 nM (magenta), 100 nM (cyan), 1 μM (blue) and 10 μM (green) #63 prior to RNase treatment. NsEMs of the NCP reassemblies are shown, colour-coded as in the LS traces. Scale bars throughout = 50 nm. NsEMs of nuclease treated samples can be found in Sup Fig 5.

**Table 4:** Titration of #63 to illustrate its effect pgRNA mediated NCP reassembly in vitro. Left to Right: A_260_ (before and after nuclease treatment), A_260/280_ ratios, packaged RNA yield, and R_*h*_ values for reassembled materials resulting from the interaction between Cp and pgRNA, in the presence of 1, 10, 100, 1000 and 10000 nM #63. Values are shown after nuclease treatment unless stated.

| Compound #63 / nM | NCP Peak |  |  |  |  |
| --- | --- | --- | --- | --- | --- |
|  | A<sub>260</sub> (Before- → After Nuclease treatment) | A<sub>260/280</sub> | RNA yield | R<sub>h</sub> / nm | NCP appearance |
| 1 | 0.031 → 0.028 | 1.49 | 90% | 17.7 | Separate NCPs, some stain excluding |
| 10 | 0.028 → 0.013 | 1.46 | 46% | 18.3 | Malformed but separate NCPs |
| 100 | 0.027 → 0.004 | 1.03 | 15% | 16.4 | Partially formed NCPs |
| 1000 | 0.024 → 0.003 | 0.86 | 8% | 18.1 | Aggregated partially formed particles |
| 10000 | 0.026 → 0.004 | 0.93 | 15% | 18.6 | Aggregated partially formed particles |

### Structure Activity Relationship (SAR) of Compound 63 and its binding site

PS-binding ligands identified in the SMM array possess varied chemophores. In order to understand structural features of Compound #63 which contribute to PS binding, and by implication an assembly inhibitor, a simple structure-activity relationship (SAR) of its pendant functional groups was carried out (Fig 5). Five commercial variants were used, mostly exploring the roles of the halogens in the C6 ring, with one variant deleting the amino group on the C5 ring. There is a >4000-fold drop in affinity for PS1 upon removal of the amino group, highlighting it as a key feature for RNA recognition. Substitutions at the halogen positions can be compensated by having an aldehyde instead of chorine, but all other combinations, including simply fluorine replacing the chlorine cause severe loss of RNA affinity. Binding of these variants to JQ707375.1 PS1 lacking a bulged stem (PS1ΔBulge) seem largely indifferent to variations of the halogens but moderately sensitive to the presence of the amino group, hinting at the orientations of those groups on the RNA. In contrast, there are mixed deleterious effects on affinity for substitutions of the halogens with a PS1 missing the –RGAG-motif (PS1ΔMotif).

**Figure 5:**
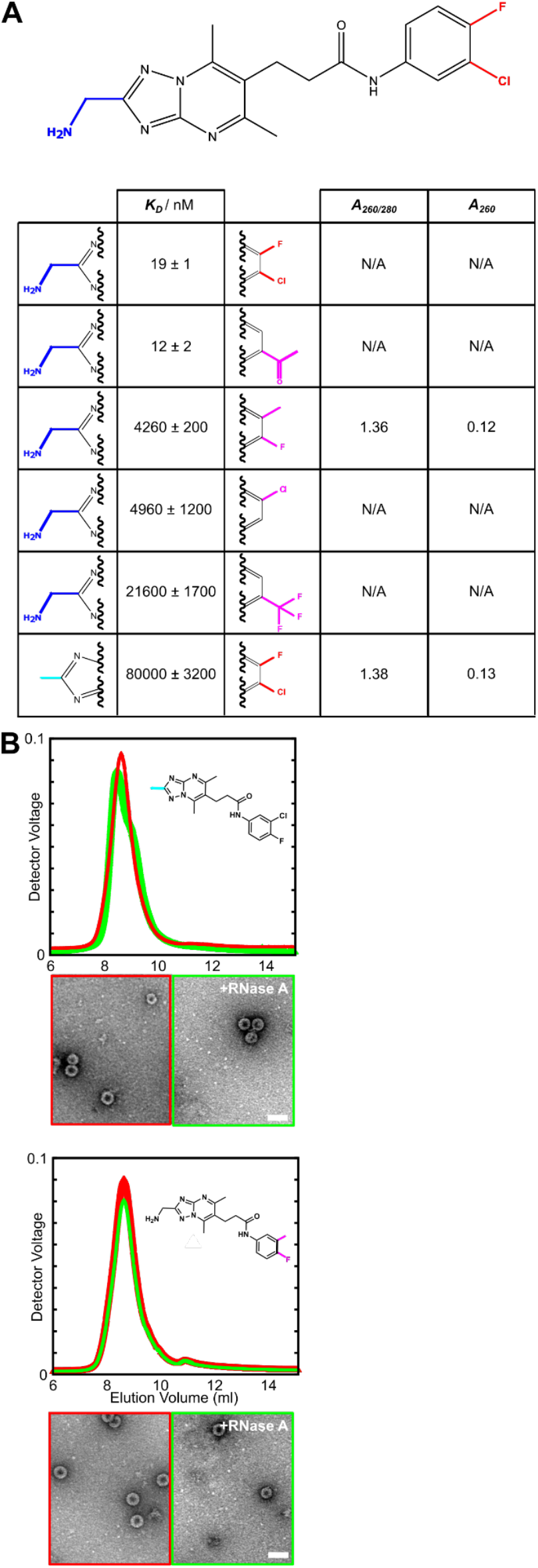
Structure-Activity Relationship investigation with #63. (A) The distal (blue and red) groups of #63 (top) were varied (cyan and magenta), generating a series of homologues. SPR was used to test the affinities of the resulting compounds for PS1 (shown here) and its two variants, PS1ΔBulge and PS1ΔMotif (Sup Fig 6). The groups varied and their corresponding affinities (*K*_*D*_) are shown in a table. The compounds which were used in NCP reassemblies are listed with their associated A_260/280_ ratios and A_260_ values. (B) *Above*, LS trace of the NCP reassemblies (performed as in Methods and as described previously^25^) between 1nM pgRNA and Cp dimer in the presence of 10 µM of inset compound -(red)/+(green) nuclease treatment. *Lower*, nsEMs of the NCP reassembly mixes colour-coded as shown in the LS trace above. Scale bars throughout = 50 nm.

### Modelling the Effects of PS-binding Compounds

Given the pressing clinical need for inhibitory reagents, we therefore sought to explore the effects of the PS-binding compounds using a mathematical model. The reaction kinetics of CP assembly around gRNA was modelled in the context of the Gillespie approach^42^, mimicking the concentrations and timings of CP and compound addition used in the experiment (see Methods). Fig. 6 shows the number of intact particles formed over time for the drug free case (black), and for different concentrations of compound 63 (1 nM, blue; 5 nM, red; 10 nM magenta). The total number of complete particles at 160 minutes, corresponding to the time particles numbers were measured experimentally, is 7.2 × 10^10^ in the drug-free case, and 1.4 × 10^10^, 1.5 × 10^7^, and 6.4 × 10^3^in the presence of 1 nM, 5nM, and 10 nM of compound 63, respectively.

**Figure 6:**
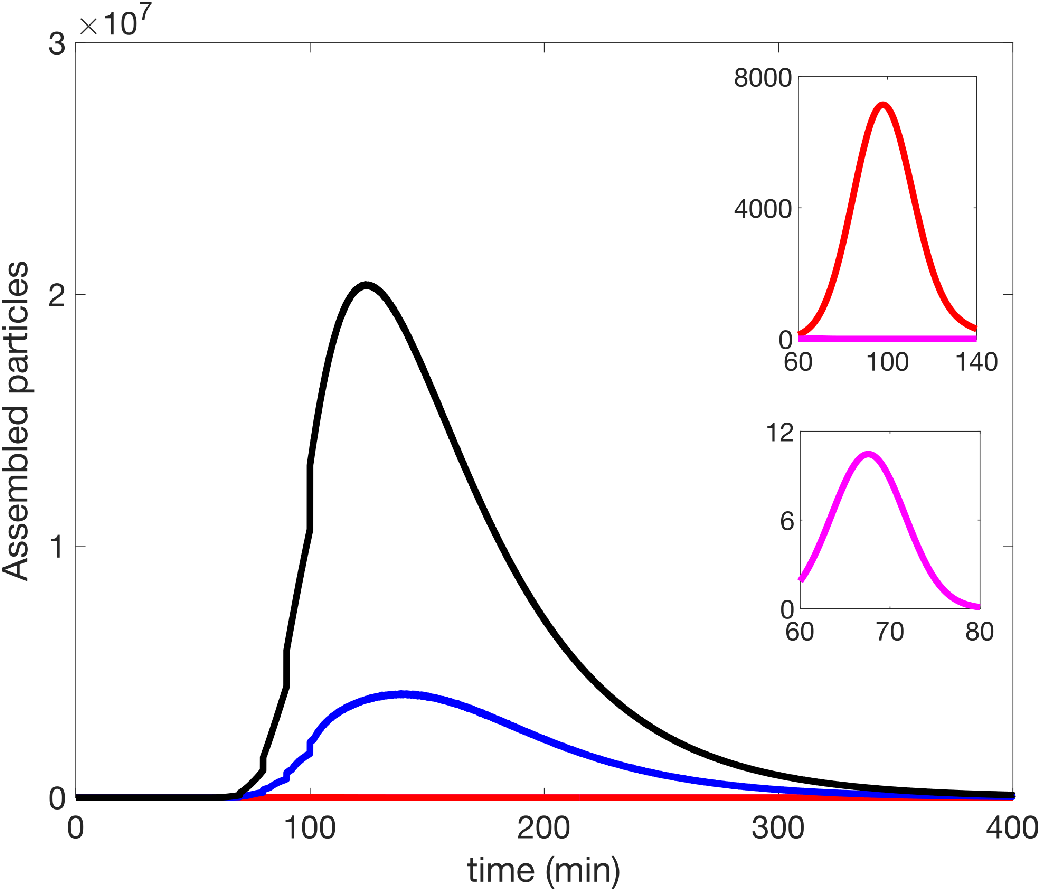
Particle assembly in the presence of different concentrations of compound 63. Particle assembly in the drug-free case (black) compared with assembly in the presence of 1 nM (blue), 5 nM (red) and 10 nM (magenta) of compound 63, respectively.

A comparative analysis with the other compounds (Table 5) was performed for 1 nM drug concentration, as well as for 3 nM, which corresponds to the stoichiometry at which there are three drug molecules per gRNA, i.e., one for each PS on each gRNA. A 10-fold increase to 30 nM, when compound 63 has ablated particle formation completely, is also shown. For 10000 nM concentration, only malformed or partially formed particles are observed (Table 3).

**Table 5:** Comparative analysis of compounds #3, #10, #46 and #63 using Intracellular Modelling. The total number of complete particles at 160 min post the start of the experiment is shown in the presence of different concentrations of each compound.

| Concentration<br>Compound | 1 nM | 3 nM | 30 nM |
| --- | --- | --- | --- |
| 3 | $7.1 \times 10^{10}$ | $6.9 \times 10^{10}$ | $5.1 \times 10^{10}$ |
| 10 | $6.2 \times 10^{10}$ | $4.7 \times 10^{10}$ | $1.0 \times 10^{10}$ |
| 46 | $6.6 \times 10^{10}$ | $5.8 \times 10^{10}$ | $2.1 \times 10^{10}$ |
| 63 | $1.4 \times 10^{10}$ | $6.0 \times 10^8$ | 0.0 |

## Discussion

There are currently very few drugs that target RNA therapeutically, although risdiplam, a small molecule targeting RNA in spinal muscular dystrophy, has recently gained FDA approval^43^. This is an important milestone since the human transcriptome is considerably larger than the proteome where until now most drug targets have been utilized. Targeting RNA with small ligands that are not based on oligonucleotides is also advantageous because of their ease of administration. Here we present data that suggest it will be possible to target viral assembly via its PS-mediated assembly. This is potentially very useful because many viral families in which we have demonstrated PS-mediated assembly^16–22^ lack effective vaccines or directly-acting drugs (DAAs).

This is the first example of a DAA targeting PS-mediated virion assembly to be identified. RNA PSs contribute collectively to make assembly efficient. They vary around a consensus sequence/motif, but they all share a common feature imbuing them with the ability to recognise their cognate CPs. Selecting ligands against HBV PS1 seems to have biased the outcome in this case such that the Cp recognition motif –RGAG-is part of the ligand binding feature. This outcome could be improved further by screening libraries with oligonucleotides encompassing several PS sites. The collective nature of PS sites ensures that they remain stable through viral evolutionary steps, and would facilitate isolation of DAAs able to work across strain variants. This seems like a promising route to isolate novel anti-viral agents.

The devastating death toll from HBV highlights the urgent need for new more effective treatment options^1,44^. The WHO has declared making HBV infection treatable by 2030 a global challenge^45^. New DAA treatments are being developed, such as Cp allostery modulators (CPAMs), which disrupt HBV nucleocapsid (NC) assembly, based on the initial discovery of the HAP compounds^46–50^. These ligands have shown early promise in clinical trials^48,51–54^. As with HIV, it may require development of drugs targeting multiple differing steps of the viral lifecycle to achieve a cure. The RNA PS-directed ligands described here dysregulate NC assembly *in vitro*, which *in vivo* might have the advantage of triggering a better immune response, and in any event would complement the action of CPAMs.

The RNA PS-directed ligands identified in the SMM screen were initially tested in *in vivo* cell culture assays of HBV replication, and several including the ones highlighted here showed partial inhibition, with the exception of Compound #3 (Dorner *pers comm*.). Unfortunately, these results have proved difficult to replicate (Tavis *pers comm*.). Therefore chemical modification to improve the bioavailability of these ligands in hepatocytes where HBV replicates will be required to achieve the goal of targeting a unique aspect of the viral lifecycle. In order to facilitate assays of such variants we established a semi-high-throughput anisotropy assembly assay, the data from which suggests we can readily identify assembly inhibitors (Sup Fig 7). The assays described throughout may assist in ligand development by allowing high-throughput screening beyond the SMM used here.

## Acknowledgements

This paper is dedicated to the memory of our colleague Prof Marcus Dorner, Imperial College, London, who passed away in 2019.

We thank Prof. Adam Zlotnick, Indiana University for gifts of his HBV Cp expression clone and insights into the production of soluble Cp for these studies. We also thank Professor John Tavis, Pennsylvania State University, for attempts to assay inhibitory effects of these PS-binding ligands *in vivo* (cell culture). We thank the Medical Research Foundation for the award of a career development grant to NP, and the UK MRC for previous grant funding to study HBV assembly (MRF-044-0002-RG-PATEL & MR/N021517/1, respectively). RT & PGS thank The Wellcome Trust (Joint Investigator Award Nos. 110145 & 110146 to PGS & RT, respectively) for funding. We acknowledge the financial support of The Trust of infrastructure and equipment in the Astbury Centre, University of Leeds (grants 089311/Z/09/Z; 090932/Z/09/Z & 106692), and the support of the Astbury Biostructure Facility by the University of Leeds. RT acknowledges additional funding via an EPSRC Established Career Fellowship (EP/R023204/1) and a Royal Society Wolfson Fellowship (RSWF\R1\180009). JS, FA and S Le G were supported by the Intramural Research Program of the National Cancer Institute, National Institutes of Health, Department of Health and Human Services.

## Supplementary Information

**Sup. Figure 1:**
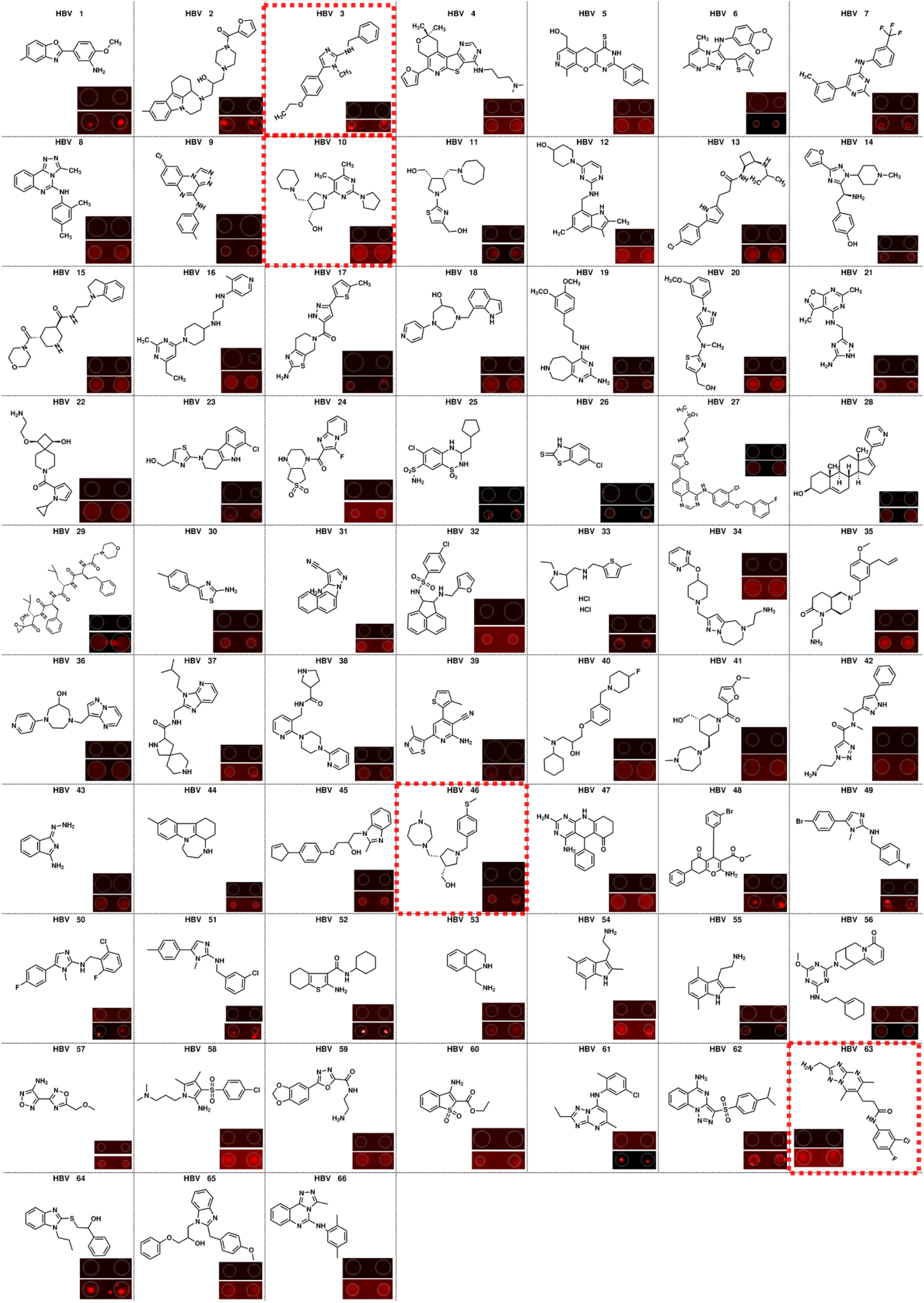
Compounds identified by SMM screen. 20,000 RNA binding compounds were screened against Cy5 labelled PS1. 72 binders were initially identified, and the 66 commercially available compounds shown here were purchased for characterisation. Compounds used throughout the study are boxed red. Inset for each compound, fluorescence on irradiation post wash (upper: buffer wash, lower: Cy5-PS1 wash).

**Sup. Figure 2:**
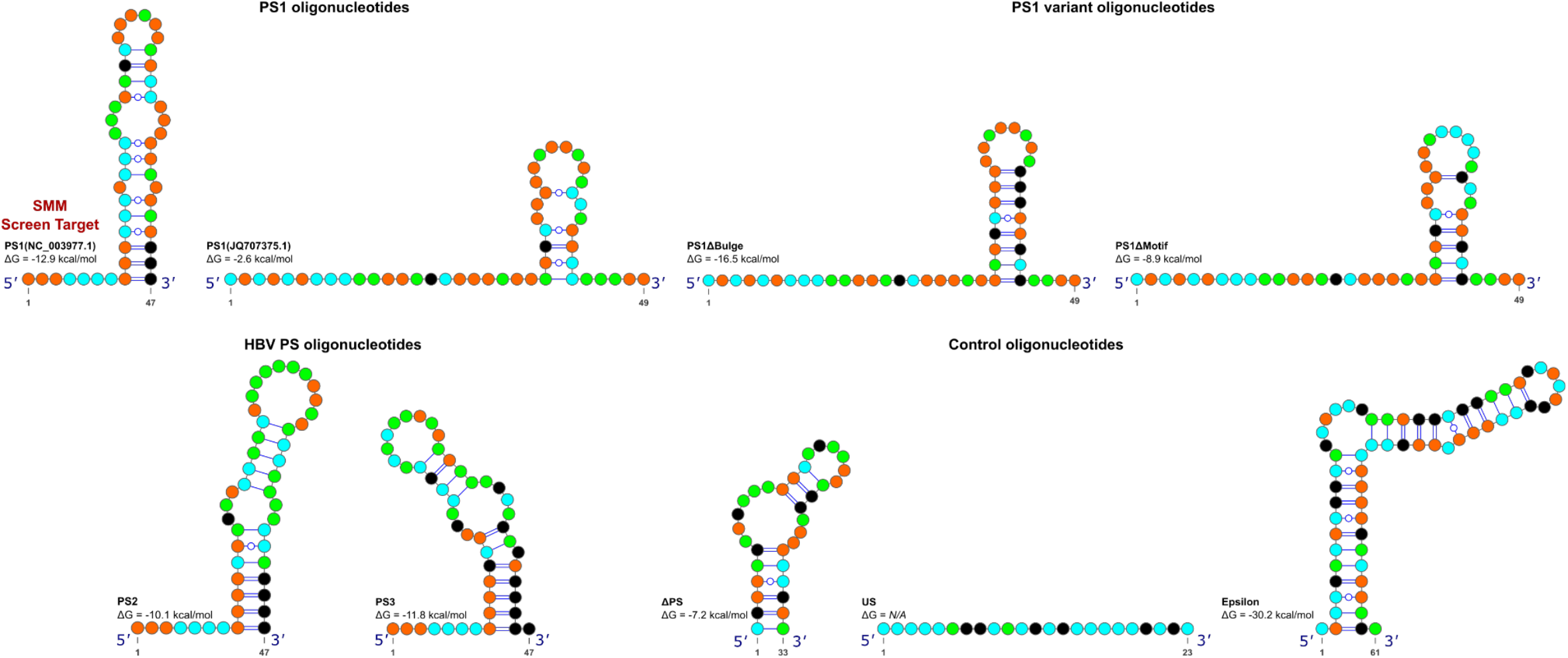
Full sequences, structures and free folding energy of RNA oligonucleotides used throughout the study. *Top*, PS1 (NC003977.1 and JQ707375.1) and variant PS1 stem loops PSΔBulge and PSΔMotif used in the SPR assays. G:C clamps were used to force the presentation of the bulged structure in the sequences from the NC_003977.1 strain, presenting the RGAG motif on an apical loop. *Bottom left*, PS stem loops PS2 and 3 from strain NC_003977.1 and *right*, control RNA oligonucleotides, ΔPS (RNA stem loop from unrelated picornavirus), US, unstructured 23-mer and epsilon, the stem loop presented at the 5′ end of the HBV pgRNA (Light green=A, orange=G, black=C, cyan=U). Free folding energy for all RNA oligomers are shown. Watson-Crick base pairs for RNA oligomers are indicated as lines, which are interrupted by circles for G-U pairs.

**Sup. Figure 3:**
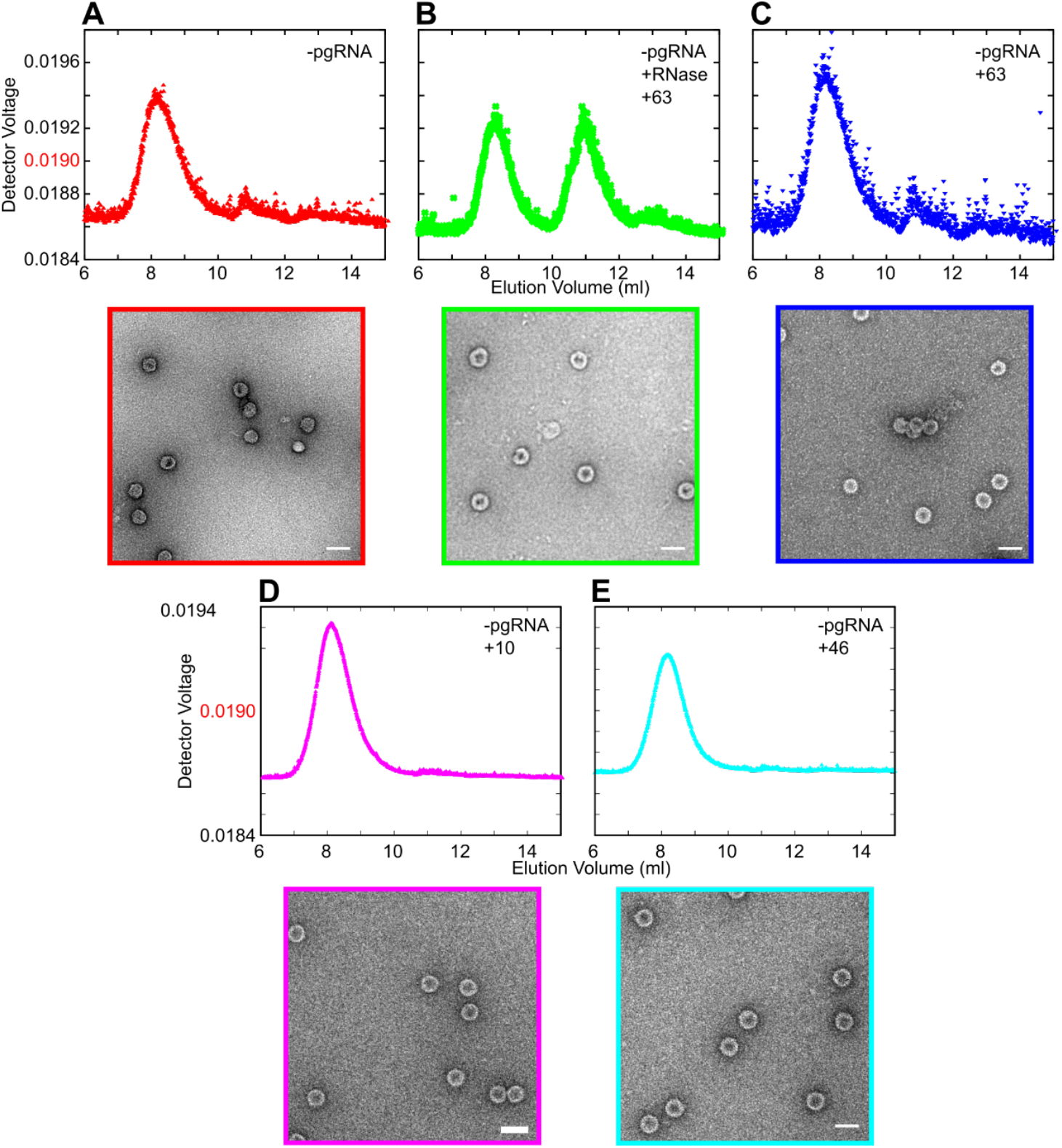
The self-association of core protein dimer is not affected by the presence of PS binding compounds. *Above*, LS traces of (A) the self-assembly of Cp dimer, (B) 10 μM #63 and 1 μM RNase A, (C) 10 μM #63, (D) 10 μM #10 and (E) 10 μM #46. R_*h*_ values taken at the arrows above each peak. *Lower*, resulting nsEM of assembly mixes, colour coded as in LS traces. Scale bars throughout = 50 nm.

**Sup. Figure 4:**
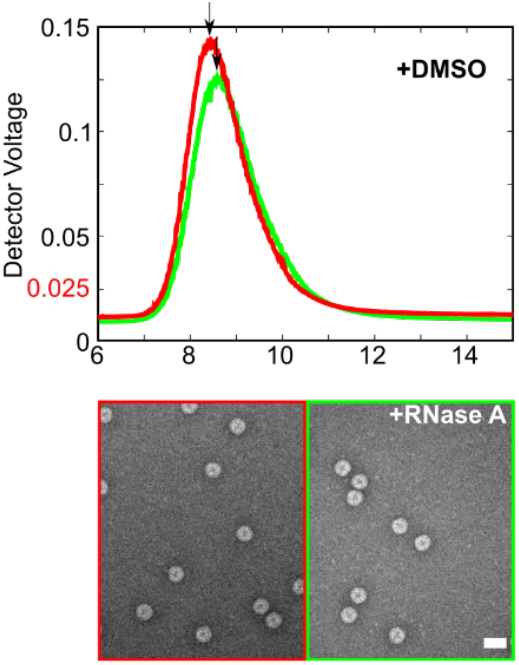
Reassembly assays to determine the effect of PS binding compounds on NCP reassembly. µM Cp dimer was titrated in gradually increasing increments into 1 nM heat annealed (Methods) pgRNA. These samples were characterized as previously described^25^, using a SEC-MALLs system and nsEM. NCP reassembly between 1 nM HBV pgRNA and HBV Cp dimer in the presence of DMSO. R_*h*_ values were taken at the arrows above each peak. *Above*, light scattering (LS) traces of the reassembly mixes, coloured-coded as described above. *Lower*, nsEMs of reassembly mixes colour-coded as in LS traces. Scale bars throughout = 50 nm.

**Sup. Figure 5:**
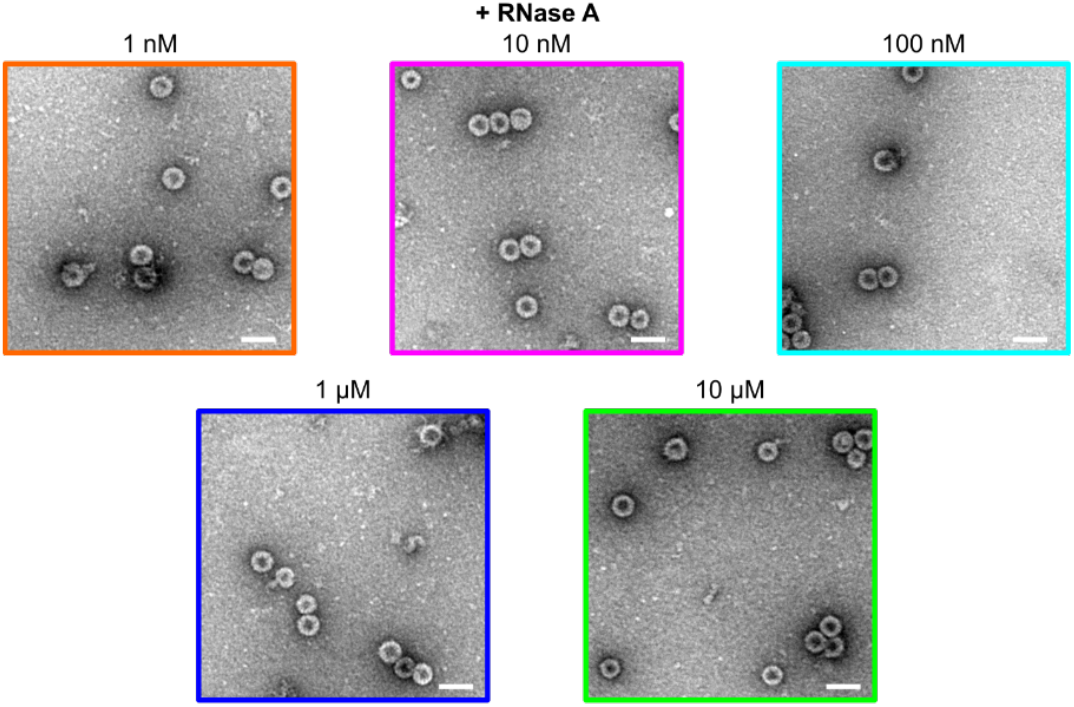
Investigating the limits of inhibition by # 63. NsEMs of the NCP reassemblies performed between 1 nM pgRNA and 1.2 μM Cp dimer in the presence of 1 nM (orange), 10 nM (magenta), 100 nM (cyan), 1 μM (blue) and 10 μM (light green) after nuclease treatment. Scale bars throughout = 50 nm.

**Sup. Figure 6:**
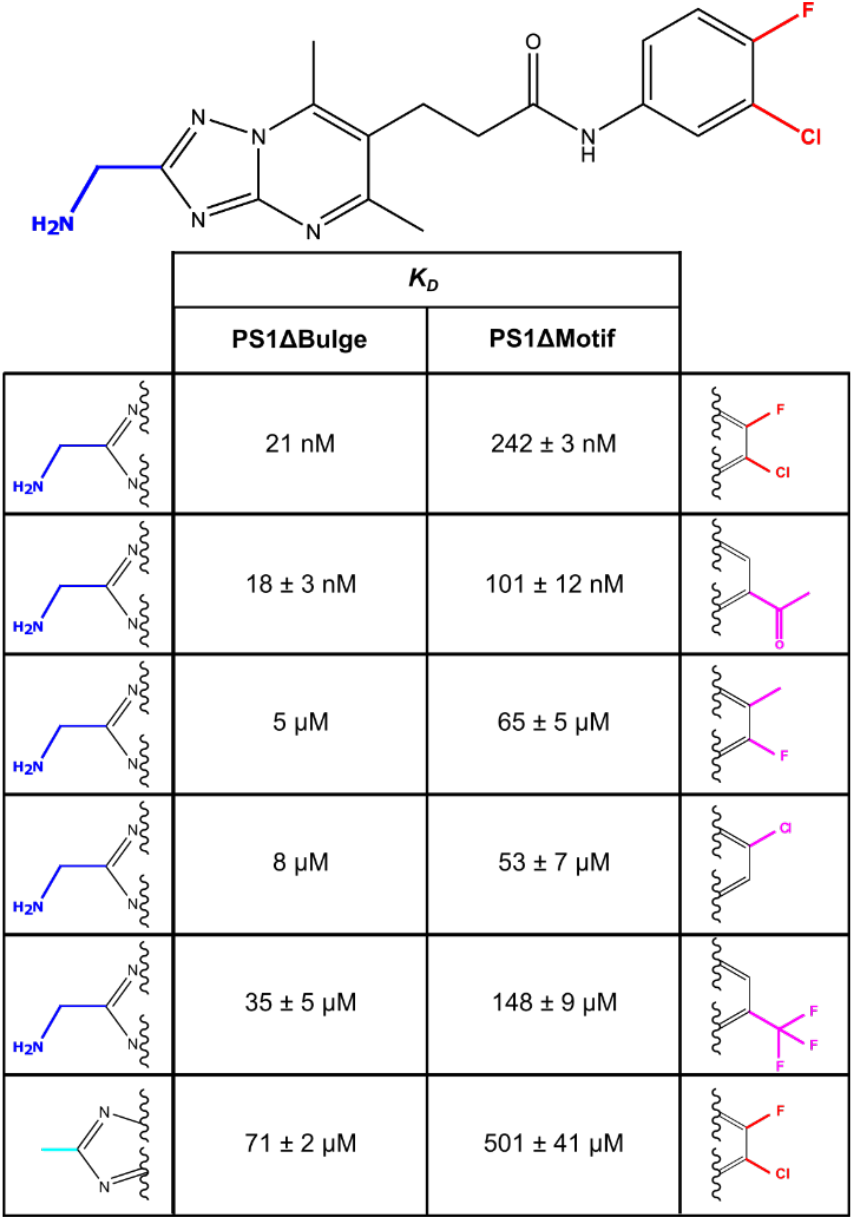
Structure-Activity Relationship investigation with #63. The distal (blue and red) groups of #63 (top) were varied (cyan and magenta), generating a series of homologous compounds, and SPR was used to test the affinities of the resulting compounds for two PS1 variants, PS1ΔBulge and PS1ΔMotif. The groups varied and their corresponding affinities (*K*_*D*_) are shown in a table (below). Errors < 1 µM are omitted for clarity.

**Sup. Figure 7:**
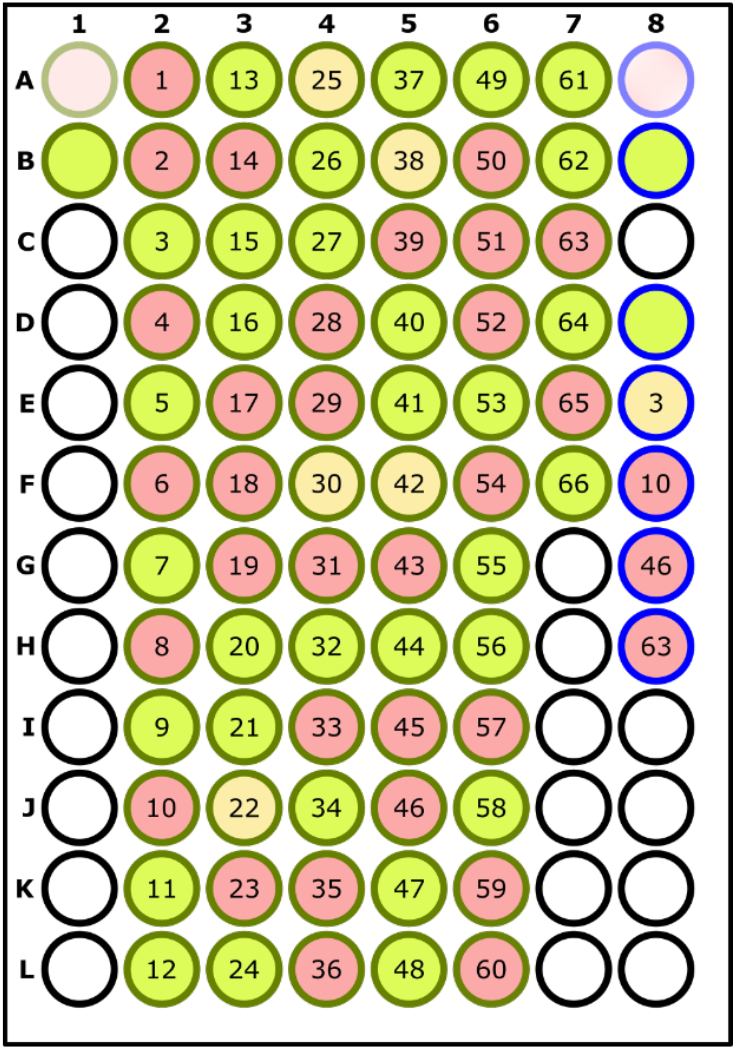
96-well plate based fluorescence anisotropy NCP assembly assay: Heat annealed 1 nM pgRNA or 15 nM PS1 oligo (NC_003977.1, Sup Fig 2) were dispensed into a 96-well plate (blue, green outlines respectively). 10 µM compound were added to wells as labelled, with an equivalent volume of DMSO added to the others. 1.2 µM Cp dimer was titrated in gradually increasing increments into the nucleic acid, as described previously (Methods) and any assembled material then challenged with 1 µM RNase A. The range of the final normalised fluorescence anisotropy change, as compared to wells A1 and A8, to which no Cp dimer is added, is denoted by the colour fill in each well. *Light red* = 0-0.3, suggestive of poor Cp dimer: RNA binding; *light yellow* = 0.3-0.6, suggestive of some Cp dimer:RNA binding; and *light green* 0.6-1, suggestive of efficient NCP assembly.

**Sup. Table 1:** Binding affinities (nM) of PS binding compounds for RNA oligonucleotides PS1, PS2 and PS3. Left to right: *Binding affinities (nM) and associated standard error of the mean of the interaction between* PS1(NC_003977.1)/PS1(JQ707375.1)/PS2/PS3 RNA oligonucleotides and compounds identified in the SMM. The standard errors of the mean *(errors <1 are omitted)* are a result of SPR measurements of 5 different compound concentrations, performed in triplicate. Control oligonucleotides show no significant binding on assayed compounds and have therefore been omitted for clarity. Where binding to controls was detected, affinities were several orders of magnitude higher than seen for PSs 1-3.

| Compound | Affinities / nM |  |  |  |
| --- | --- | --- | --- | --- |
|  | PS1 (NC 003977.1) | PS1 (JQ707375.1) | PS2 | PS3 |
| 1 | 165 ± 2 |  | 141 ± 4 | 1755 ± 125 |
| 2 | 10027 ± 119 |  | 86441 ± 101 | 44115 ± 28 |
| 3 | 211982 ± 1179 | 224056 ± 455 | 320 ± 5 | 99060 ± 591 |
| 4 | 171800 ± 24900 |  | 1913 ± 83 | 59209 ± 3220 |
| 5 | 239 ± 1 |  | 50359 ± 1890 | 312 ± 1 |
| 6 | 61 ± 2 |  | 112 ± 1 | 615 ± 22 |
| 7 | 456 ± 5 |  | 174364 ± 248 | 491 ± 5 |
| 8 | 31 ± 2 | 92 ± 12 | 385281 ± 1040 | 1722 ± 1 |
| 9 | 30 ± 10 |  | 2891 ± 45 | 37 ± 3 |
| 10 | 11773 ± 284 | 11374 ± 169 | 10899 ± 691 | 8539 ± 797 |
| 11 | 11498 ± 220 |  | 147 ± 35 | 27121 ± 1234 |
| 12 | 61859 ± 4538 |  | 46304 ± 271 | 94712 ± 8466 |
| 13 | 20114 ± 20 |  | 26209 ± 375 | 206 ± 17 |
| 14 | 345761 ± 22 |  | 65111 ± 167 | 161091 ± 1010 |
| 15 | 32 ± 3 |  | 6 ± 4 | 1 ± 2 |
| 16 | 18364 ± 92 |  | 22984 ± 135 | 21547 ± 159 |
| 17 | 135 | 421 ± 54 | 199135 ± 402 | 9964365 ± 2000 |
| 18 | 83580 ± 700 |  | 124123 ± 5250 | 58575 ± 2180 |
| 19 | 19336 ± 900 | 12164 ± 231 | 16890 ± 565 | 16820 ± 1140 |
| 20 | 11 ± 5 |  | 66012 ± 3479 | 2286 ± 23 |
| 21 | 20 ± 2 |  | 30379 ± 79 | 3934 ± 526 |
| 22 | 18551 ± 264 |  | 37839 ± 29 | 932724 ± 4200 |
| 23 | 429159 ± 50 |  | 31 ± 6 | 368 ± 3 |
| 24 | 15 ± 4 |  | 14364 ± 20 | 44 ± 7 |
| 25 | 732 ± 57 | 1241 ± 87 | 42202 ± 29 | 12 ± 4 |
| 26 | 66 ± 9 |  | 2764 ± 2 | 52 |
| 27 | 816 ± 12 |  | 3961266 ± 7012 | 50769585 ± 1330 |
| 28 | 883 ± 4 |  | 57 ± 3 | 677 ± 8 |
| 29 | 16 ± 8 | 28 ± 9 | 182 ± 2 | 73 ± 17 |
| 30 | 177 |  | 169220 ± 8600 | 11 ± 1 |
| 31 | 154799 ± 6999 |  | 65457 ± 191 | 18 ± 1 |
| 32 | 64347 ± 32 |  | 547775 ± 4680 | 36 |
| 33 | 751168 ± 186 |  | 113994 ± 1950 | 1756 |
| 34 | 374800 ± 1500 |  | 116771 ± 3280 | 60460 ± 1600 |
| 35 | 1221 ± 150 | 1943 ± 65 | 348 ± 51 | 4739 ± 378 |
| 36 | 585 ± 37 |  | 239 ± 2 | 358037 ± 1000 |
| 37 | 37 ± 2 |  | 258296 ± 1350 | 3748 ± 3 |
| 38 | 55 ± 2 | 184 ± 31 | 124 | 1 ± 5 |
| 39 | 3 ± 7 |  | 467 ± 2 | 4543 ± 328 |
| 40 | 832300 ± 2750 |  | 58889 ± 7090 | 22725 ± 2500 |
| 41 | 532 ± 12 |  | 241834 ± 1380 | 697548 ± 53 |
| 42 | 18 ± 1 |  | 2094 ± 1 | 388 ± 32 |
| 43 | 17 ± 1 |  | 74 | 19 ± 1 |
| 44 | 519686 ± 546 |  | 173 ± 27 | 589734 ± 14600 |
| 45 | 248562 ± 371 |  | 17176 ± 798 | 162 ± 3 |
| 46 | 10114 ± 119 | 10195 ± 224 | 16583 ± 336 | 32657 ± 227 |
| 47 | 2 |  | 99 ± 2 | 73138 ± 4740 |
| 48 | 384 ± 5 |  | 112908 ± 572 | 90 ± 7 |
| 49 | 256 ± 24 |  | 83838 ± 1840 | 311148 ± 432 |
| 50 | 7 ± 3 |  | 170 ± 2 | 1246780 ± 850 |
| 51 | 133 ± 9 |  | 403424 ± 63500 | 336347 ± 100 |
| 52 | 7 ± 6 |  | 4661090 ± 7960 | 9 ± 2 |
| 53 | 751 ± 12 | 621 ± 56 | 299320 ± 7970 | 935 ± 10 |
| 54 | 37980 ± 189 |  | 7123 ± 1970 | 3599335 ± 26 |
| 55 | 24281 ± 1071 |  | 456639 ± 7680 | 1161100 ± 3233 |
| 56 | 1312 ± 3 | 768 ± 98 | 11 ± 2 | 57 ± 7 |
| 57 | 9 ± 5 |  | 227 ± 1 | 20 ± 11 |
| 58 | 23424 ± 243 |  | 13654 ± 8190 | 24618 ± 2230 |
| 59 | 0.1 ± 2 |  | 69 ± 1 | 29 ± 3 |
| 60 | 5 ± 3 |  | 49 ± 5 | 59320 ± 620 |
| 61 | 2 ± 4 |  | 33999 ± 232 | 217693 ± 1400 |
| 62 | 2002 ± 234 |  | 516618 ± 22700 | 4617 ± 392 |
| 63 | 12 | 19 | 20 ± 1 | 14 |
| 64 | 10 ± 7 |  | 46 | 79 ± 3 |
| 65 | 14 |  | 1 | 28 ± 4 |
| 66 | 272 ± 6 |  | 100 ± 1 | 2 |

**Table 2:** Binding affinities (nM) and associated standard error of the mean (errors <1 are omitted) of PS binding compounds used throughout, for RNA oligonucleotides PS(JQ707375.1) and its variants, PS1ΔMotif and PS1ΔBulge.

| Compound | $K_D$ / nM | | |
| --- | --- | --- | --- |
| | PS1(JQ707375.1) | PS1ΔMotif | PS1ΔBulge |
| 3 | 224056 ± 455 | 421401 ± 367 | 312051 ± 391 |
| 10 | 11374 ± 169 | 48245 ± 258 | 10397 ± 525 |
| 46 | 10196 ± 225 | 35548 ± 126 | 9658 ± 227 |
| 63 | 19 | 242 ± 3 | 21 |

**Table 3:**
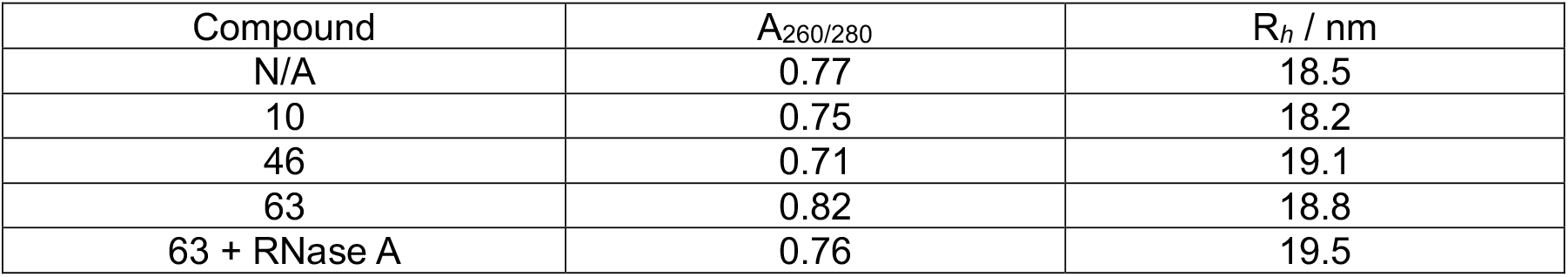
Effect of Compounds on self-assembly of Cp dimers into NCPs in vitro. Left to Right: A_260/280_ ratios and R_*h*_ values for reassembled materials resulting from the self-assembly of Cp dimers, in the absence and presence of compounds 10, 46, and 63.

## References

1. Mathers, B. M. et al.. Global epidemiology of injecting drug use and HIV among people who inject drugs: a systematic review. Lancet 372, 1733–1745 (2008).

2. Murray, K. et al.. Protective immunisation against hepatitis B with an internal antigen of the virus. J. Med. Virol. (1987) doi:10.1002/jmv.1890230202.

3. Tillmann, H. L. Antiviral therapy and resistance with hepatitis B virus infection. World J. Gastroenterol. (2007) doi:10.3748/wjg.v13.i1.125.

4. Lok, A. S. et al.. Long-term safety of lamivudine treatment in patients with chronic hepatitis B. Gastroenterology 125, 1714–1722 (2003).

5. Seeger, C. & Mason, W. S. Hepatitis B virus biology. Microbiol Mol Biol Rev 64, 51–68 (2000).

6. Bartenschlager, R. & Schaller, H. Hepadnaviral assembly is initiated by polymerase binding to the encapsidation signal in the viral RNA genome. EMBO J 11, 3413–3420 (1992).

7. Bartenschlager, R., Junker-Niepmann, M. & Schaller, H. The P gene product of hepatitis B virus is required as a structural component for genomic RNA encapsidation. J. Virol. (1990) doi:10.1128/jvi.64.11.5324-5332.1990.

8. Junker-Niepmann, M., Bartenschlager, R. & Schaller, H. A short cis-acting sequence is required for hepatitis B virus pregenome encapsidation and sufficient for packaging of foreign RNA. EMBO J. (1990) doi:10.1002/j.1460-2075.1990.tb07540.x.

9. Selzer, L. & Zlotnick, A. Assembly and Release of Hepatitis B Virus. Cold Spring Harb Perspect Med 5, (2015).

10. Beck, J. & Nassal, M. Hepatitis B virus replication. World J Gastroenterol 13, 48–64 (2007).

11. Nassal, M. Hepatitis B viruses: Reverse transcription a different way. Virus Res. (2008) doi:10.1016/j.virusres.2007.12.024.

12. Wang, J. C., Nickens, D. G., Lentz, T. B., Loeb, D. D. & Zlotnick, A. Encapsidated hepatitis B virus reverse transcriptase is poised on an ordered RNA lattice. Proc Natl Acad Sci U S A 111, 11329–11334 (2014).

13. Wei, L. & Ploss, A. Hepatitis B virus cccDNA is formed through distinct repair processes of each strand. Nat. Commun. 12, (2021).

14. Guo, Y. H., Li, Y. N., Zhao, J. R., Zhang, J. & Yan, Z. HBc binds to the CpG islands of HBV cccDNA and promotes an epigenetic permissive state. Epigenetics 6, 720–726 (2011).

15. Rolfsson, O. et al.. Direct Evidence for Packaging Signal-Mediated Assembly of Bacteriophage MS2. J Mol Biol 428, 431–448 (2016).

16. Shakeel, S. et al.. Genomic RNA folding mediates assembly of human parechovirus. Nat Commun 8, 5 (2017).

17. Chandler-Bostock, R. et al.. Assembly of infectious enteroviruses depends on multiple, conserved genomic RNA-coat protein contacts. PLoS Pathog. 16, (2020).

18. Tetter, S. et al.. Evolution of a virus-like architecture and packaging mechanism in a repurposed bacterial protein. Science (80-.). 372, 1220–1224 (2021).

19. Patel, N. et al.. HBV RNA pre-genome encodes specific motifs that mediate interactions with the viral core protein that promote nucleocapsid assembly. Nat. Microbiol. 2, 17098 (2017).

20. Borodavka, A., Tuma, R. & Stockley, P. G. A two-stage mechanism of viral RNA compaction revealed by single molecule fluorescence. RNA Biol 10, 481–489 (2013).

21. Twarock, R. & Stockley, P. G. RNA-Mediated Virus Assembly: Mechanisms and Consequences for Viral Evolution and Therapy. Annual Review of Biophysics (2019) doi:10.1146/annurev-biophys-052118-115611.

22. Bunka, D. H. et al.. Degenerate RNA packaging signals in the genome of Satellite Tobacco Necrosis Virus: implications for the assembly of a T=1 capsid. J Mol Biol 413, 51–65 (2011).

23. Dykeman, E. C., Stockley, P. G. & Twarock, R. Solving a Levinthal’s paradox for virus assembly identifies a unique antiviral strategy. Proc Natl Acad Sci U S A 111, 5361–5366 (2014).

24. Bunka, D. H. J. & Stockley, P. G. Aptamers come of age - At last. Nature Reviews Microbiology (2006) doi:10.1038/nrmicro1458.

25. Patel, N. et al.. Functional Analysis of Essential pgRNA Sites Regulating Assembly of Hepatitis B Virus (Under Review). Commun. Biol.

26. Connelly, C. M., Abulwerdi, F. A. & Schneekloth, J. S. Discovery of RNA binding small molecules using small molecule microarrays. in Methods in Molecular Biology (2017). doi:10.1007/978-1-4939-6584-7_11.

27. Abulwerdi, F. A. et al.. Development of Small Molecules with a Noncanonical Binding Mode to HIV-1 Trans Activation Response (TAR) RNA. J. Med. Chem. (2016) doi:10.1021/acs.jmedchem.6b01450.

28. Sztuba-Solinska, J. et al.. Identification of biologically active, HIV TAR RNA-binding small molecules using small molecule microarrays. J. Am. Chem. Soc. (2014) doi:10.1021/ja502754f.

29. Connelly, C. M., Boer, R. E., Moon, M. H., Gareiss, P. & Schneekloth, J. S. Discovery of Inhibitors of MicroRNA-21 Processing Using Small Molecule Microarrays. ACS Chem. Biol. (2017) doi:10.1021/acschembio.6b00945.

30. Felsenstein, K. M. et al.. Small Molecule Microarrays Enable the Identification of a Selective, Quadruplex-Binding Inhibitor of MYC Expression. ACS Chem. Biol. (2016) doi:10.1021/acschembio.5b00577.

31. Porterfield, J. Z. et al.. Full-length hepatitis B virus core protein packages viral and heterologous RNA with similarly high levels of cooperativity. J Virol 84, 7174–7184 (2010).

32. Porterfield, J. Z. & Zlotnick, A. A simple and general method for determining the protein and nucleic acid content of viruses by UV absorbance. Virology 407, 281–288 (2010).

33. Bradner, J. E. et al.. A Robust Small-Molecule Microarray Platform for Screening Cell Lysates. Chem. Biol. (2006) doi:10.1016/j.chembiol.2006.03.004.

34. Biacore & Guides, A. Kinetics and affinity measurements with Biacore TM systems.

35. Karlssonz, R. & Fält, A. Experimental design for kinetic analysis of protein-protein interactions with surface plasmon resonance biosensors. J. Immunol. Methods (1997) doi:10.1016/S0022-1759(96)00195-0.

36. Fatehi, F. et al.. An intracellular model of hepatitis b viral infection: An in silico platform for comparing therapeutic strategies. Viruses 13, (2021).

37. Endres, D. & Zlotnick, A. Model-based analysis of assembly kinetics for virus capsids or other spherical polymers. Biophys. J. 83, 1217 (2002).

38. Uetrecht, C. et al.. High-resolution mass spectrometry of viral assemblies: molecular composition and stability of dimorphic hepatitis B virus capsids. Proc. Natl. Acad. Sci. U. S. A. 105, 9216–9220 (2008).

39. Pollard, T. D. A Guide to Simple and Informative Binding Assays. https://doi.org/10.1091/mbc.e10-08-0683 21, p4061–4067 (2017).

40. Routh, A., Domitrovic, T. & Johnson, J. E. Host RNAs, including transposons, are encapsidated by a eukaryotic single-stranded RNA virus. Proc Natl Acad Sci U S A 109, 1907–1912 (2012).

41. Thai, H. et al.. Convergence and coevolution of Hepatitis B virus drug resistance. Nat. Commun. 3, 789 (2012).

42. Gillespie, D. T. Exact stochastic simulation of coupled chemical reactions. J. Phys. Chem. 81, 2340–2361 (1977).

43. Ratni, H., Scalco, R. S. & Stephan, A. H. Risdiplam, the First Approved Small Molecule Splicing Modifier Drug as a Blueprint for Future Transformative Medicines. ACS Med. Chem. Lett. (2021) doi:10.1021/ACSMEDCHEMLETT.0C00659.

44. World Health Organization. Progress report on HIV, viral hepatitis and sexually transmitted infections, 2019. 1, 1–39 (2019).

45. World Health Organization. Combating hepatitis B and C to reach elimination by 2030. 1–16 (2016).

46. Stray, S. J. et al.. A heteroaryldihydropyrimidine activates and can misdirect hepatitis B virus capsid assembly. Proc. Natl. Acad. Sci. U. S. A. (2005) doi:10.1073/pnas.0409732102.

47. Sj, S., P, C. & A, Z. Zinc ions trigger conformational change and oligomerization of hepatitis B virus capsid protein. Biochemistry 43, 9989–9998 (2004).

48. Stray, S. J. & Zlotnick, A. BAY 41-4109 has multiple effects on Hepatitis B virus capsid assembly. J. Mol. Recognit. (2006) doi:10.1002/jmr.801.

49. K, D. et al. Inhibition of hepatitis B virus replication by drug-induced depletion of nucleocapsids. Science 299, 893–896 (2003).

50. Campagna, M. R. et al.. Sulfamoylbenzamide Derivatives Inhibit the Assembly of Hepatitis B Virus Nucleocapsids. J. Virol. 87, 6931 (2013).

51. Zoulim, F. et al.. Safety, tolerability, pharmacokinetics and antiviral activity of JNJ-56136379, a novel HBV capsid assembly modulator, in non-cirrhotic, treatment-naive subjects with chronic hepatitis B. Hepatology (2017) doi: http://dx.doi.org/10.1002/hep.29634.

52. H., Z. et al. Safety, pharmacokinetics and anti-viral efficacy of novel core protein allosteric modifier GLS4 in patients with chronic hepatitis B: Interim results from a 48 weeks phase 2a study. Hepatology (2018).

53. Mani, N. et al.. Preclinical profile of AB-423, an inhibitor of hepatitis B virus pregenomic RNA encapsidation. Antimicrob. Agents Chemother. (2018) doi:10.1128/AAC.00082-18.

54. Mf, Y. et al.. Antiviral Activity, Safety, and Pharmacokinetics of Capsid Assembly Modulator NVR 3-778 in Patients with Chronic HBV Infection. Gastroenterology 156, 1392-1403.e7 (2019).

